# IFITM1 inhibits Henipavirus membrane fusion by trapping ephrinB2 receptors in fusion-unfavorable membrane nanodomains

**DOI:** 10.64898/2026.05.06.723334

**Authors:** Yuhang Luo, Jingjing Wang, Jiake Li, Qian Wang, Jacqueline Sung, Vicky Kliemke, Chen Liang, Qian Liu

## Abstract

Nipah (NiV) and Hendra (HeV) viruses are prototype henipaviruses (HNV) within the family of *Paramyxoviridae*. They are highly pathogenic zoonotic paramyxoviruses that cause fatal encephalitis in humans, with case fatality rates of 40-70%. These viruses use the receptor-binding protein (G) to engage host receptors ephrinB2 and/or ephrinB3, while the fusion protein (F) drives membrane fusion leading to virus entry and cell-cell fusion. Interferon-inducible transmembrane (IFITM) proteins restrict the infection of many unrelated viruses at virus entry. We show that the knockdown of IFITM1 enhances HNV entry and cell-cell fusion in both human epithelial and endothelial cells, while overexpression of IFITM1 inhibits both. Using single-molecule localization microscopy, we observed that IFITM1 forms nanoclusters on the plasma membrane that are enhanced upon interferon stimulation. IFITM1 clusters spatially co-occur with GM1-enriched, raft-like membrane domains. Notably, IFITM1 interacts with ephrinB2, and redistributes ephrinB2 into larger and denser clusters, potentially within GM1-enriched regions. Using single-particle tracking, we further show that ephrinB2’s lateral diffusion on the plasma membrane is confined by IFITM1, and this confinement is not reversed by amphotericin B, a membrane-permeabilizing agent that increases membrane fluidity. Our data suggest that IFITM1 sequesters ephrinB2 in fusion-unfavorable membrane domains, thereby enhancing the energy barrier for virus-cell membrane fusion and decreasing the virus-receptor encounters.

**Significance:** Nipah (NiV) and Hendra (HeV) viruses are zoonotic paramyxoviruses that cause fatal encephalitis in humans. Here, we show that IFITM1 inhibits NiV and HeV entry at the surface of epithelial and endothelial cells. We report that IFITM1 restricts NiV entry by sequestering the host receptor ephrinB2 in membrane domains that are energetically unfavorable for virus-cell membrane fusion. IFITM1 also restricts ephrinB2 lateral diffusion on the plasma membrane, thereby reducing productive virus-receptor encounters. Notably, amphotericin B treatment, which increases membrane fluidity, did not rescue IFITM1-mediated confinement of ephrinB2 or restore NiV entry. Our data reveal a new mechanism by which IFITM1 restricts NiV infection, suggesting that its antiviral activity extends beyond simply altering membrane fluidity.

## Introduction

Interferons (IFNs) are antiviral cytokines that combat viral infections by stimulating the expression of interferon-stimulated genes (ISGs). Among these ISG products, interferon-inducible transmembrane proteins (IFITMs) 1, 2, and 3 are central players and belong to the CD225 protein family(1). Evidence supports that IFITMs prevent the infection of unrelated viruses by blocking virus-cell membrane fusion during the virus entry step(2). Although IFITM1, 2, and 3 proteins are highly conserved in the CD225 domains, IFITM1 differs from IFITM2 and 3 with a 21-residue deletion at the N terminus(2). This N-terminal region contains a 20YEML23 sorting signal that localizes IFITM2 and 3 to endosomes, and the lack of this region localizes IFITM1 to the cell surface(3, 4). The abundance of IFITM1 at the plasma membrane allows restriction of viruses entering at the cell surface, including herpesviruses (5), paramyxoviruses (6), and retroviruses (7, 8). Meanwhile, cumulative evidence suggests context-dependent and multifaceted roles of IFITMs in modulating virus infections (9–12).

IFITM proteins restrict viruses either by directly engaging viral components or by altering the membrane environment in which fusion occurs(1, 2). For the latter one, most mechanistic studies have been focused on IFITM3 inhibition of Influenza A virus (IAV) (4, 13–19). *In vivo* relevance of this restriction has been demonstrated using IFITM3 knockout mice and further supported by the association between the severity of IAV infections and single nucleotide polymorphisms in the IFITM3 gene of hospitalized patients in the seasonal and epidemic flu (20–22). The current model is that IFITMs inhibit viral entry by remodeling membrane biophysical properties rather than by blocking receptor binding *per se*. Evidence shows that IFITM proteins restrict viral glycoprotein-induced membrane fusion at least in part, by reducing membrane fluidity and altering membrane curvature (14, 19, 23). Consistently, IFITM3-mediated restriction of IAV is counteracted by amphotericin B, an antimycotic heptean, further indicating that IFITM-mediated virus restriction may be linked to altered membrane fluidity (24). S-palmitoylation is important for the membrane association of IFITM proteins, especially IFITM1 and IFITM3, and is critical for their antiviral activity against IAV (16). Structurally, a short amphipathic helix (AH) and a GxxxL motif in IFITM3 were identified as determinants of its antiviral activity and may contribute to this function by inducing membrane curvature and decreasing membrane fluidity, respectively (14, 25). Meanwhile, single-virus imaging studies show that IFITM3 blocks fusion-pore formation following IAV-endosome hemifusion (17). More recently, this notion was supported by a local lipid-sorting model in which IFITM3 redistributes lipids at the fusion site to stabilize nonproductive hemifusion intermediates and prevent fusion pore formation (26), highlighting that IFITM proteins may restrict virus entry, at least in part, through localized, precise remodeling of the membrane environment at individual virus-cell fusion sites.

The family of *Paramyxoviridae* contains many well-documented highly pathogenic human viruses with global health burdens (*e*.*g*. Measles virus, human parainfluenza viruses) and emerging zoonotic viruses (*e*.*g*. Langya virus) (27). Moreover, a global survey identified over 66 novel paramyxoviruses in bats and rodents, these extensively diverse viruses span more than eighty bat species sampled, pose the grave danger of cross-species spillover into humans (28). Among these, *Henipavirus* (HNV), a genus in the family of *Paramyxoviridae*, contains many highly pathogenic zoonotic viruses that pose known threats to human health, such as Nipah virus (NiV) and Hendra virus (HeV). NiV and HeV cause deadly encephalitis in humans and domestic animals. Yearly outbreaks of NiV infections have been reported in Southeast Asia, Bangladesh, and India(29). HNV entry into cells requires a receptor-binding protein (G) that binds to ephrinB2 and/or -B3 and a fusion protein (F) that executes membrane fusion at neutral pH (30). Cell-cell membrane fusion forms multinucleated cells (syncytia), representing a pathological hallmark of HeV and NiV infection (31). IFITM1 has been suggested to restrict the infection of several paramyxoviruses(6), and a recent study shows that IFITM3 promotes NiV envelope protein-mediated entry by interacting with NiV-F (32). Here, we show that IFITM1 is more potent in restricting membrane fusion induced by HNV glycoproteins, compared to IFITM2 and IFITM3. Mechanistically, IFITM1 traps ephrinB2 receptors within enlarged GM1-enriched membrane domains, thereby reducing the boundary between the liquid ordered and disordered membrane regions, which may in turn increase the energy barrier for virus-cell membrane fusion. IFITM1 also restricts the lateral diffusion of ephrinB2, thereby limiting the productive virus-receptor encounters.

## Results

### Endogenous IFITMs inhibit HNV glycoprotein-induced membrane fusion in endothelial and epithelial cells

Histopathological analyses of HNV infections revealed pronounced endothelial and neural tropism because both ephrinB2 and B3 receptors are present in endothelial cells, epithelial cells, glial cells, and neurons (33–35). We chose hTERT-immortalized human microvascular endothelial cells (HuMEC) as a model endothelial cell line and HEK293T cells as an epithelial cell line to study the role of endogenous IFITMs in the HNV entry (30). We performed siRNA knockdown of IFITMs in both HuMEC and HEK293T cells. Quantitative polymerase chain reaction analysis (qPCR) using primers specific for the three IFITM genes shows that substantial knockdown of IFITM1, 2, and 3 was achieved using respective siRNAs in untreated and IFN-α stimulated HuMEC (Fig. 1a-1c) and HEK293T (Fig. 1g-1i) cells, which was further confirmed by the data of western blot (Fig. 1d and 1j). We also noticed increased IFITMs expression in both cell lines upon IFN-α stimulation (Fig. 1d and 1j). Additionally, basal IFITM1 expression was barely detectable in both cell lines, whereas IFITM2 and IFITM3 were readily detectable without IFN-α stimulation (Fig. 1d and 1j). We confirmed that siRNA knockdown of IFITMs does not affect the viability of HEK293T and endothelial cell lines (Fig. S1a and S1b). We observed cross-knockdown of IFITM2 and IFITM3 but not IFITM1 (Fig. S1c). This is due to the high sequence homology between IFITM2 and IFITM3.

**Fig. 1.**
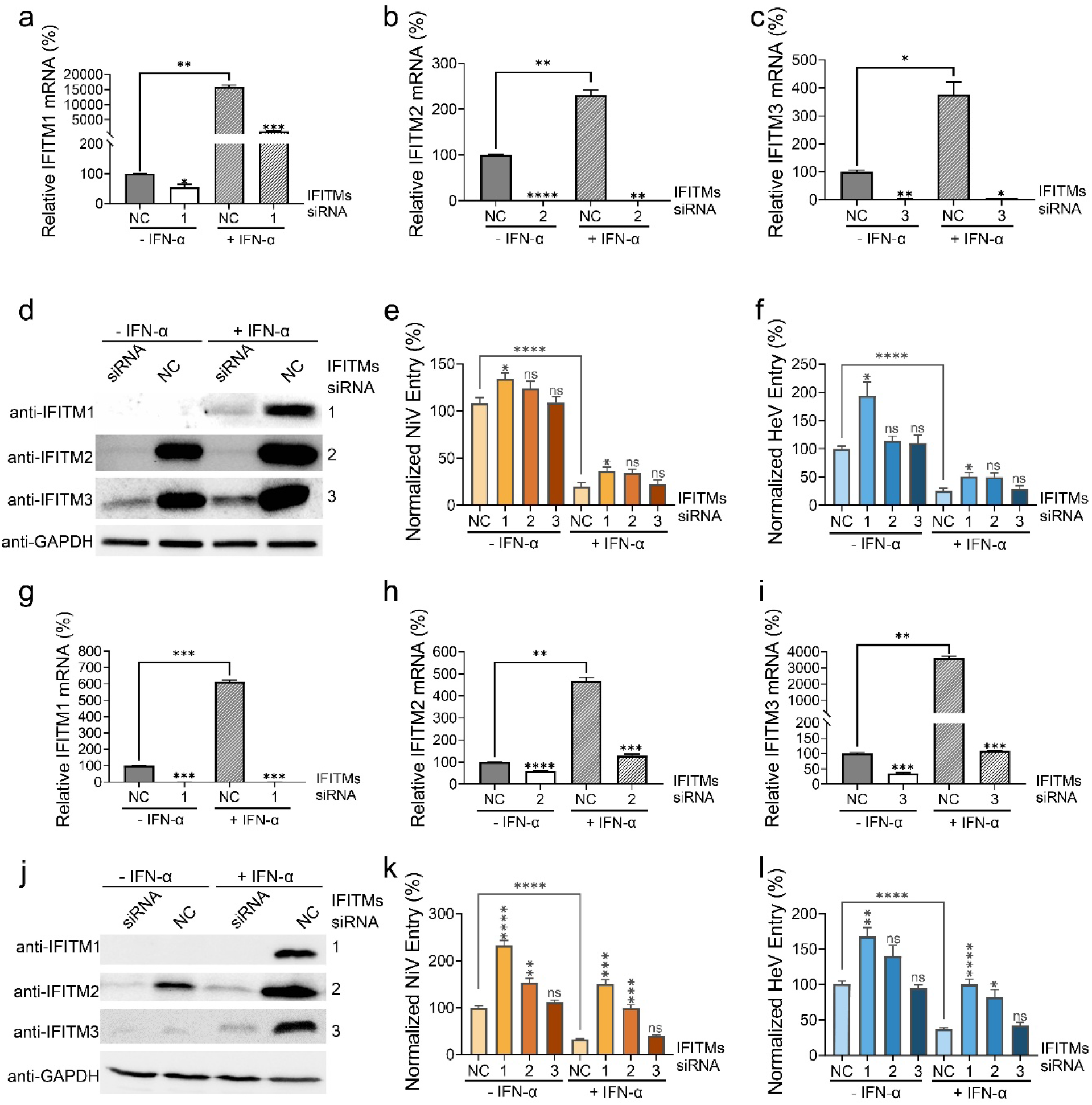
The role of endogenous IFITMs in NiV and HeV pseudovirus entry into human endothelial and epithelial cells. The endogenous IFITMs mRNA levels in HuMEC (**a-c**) and HEK293T cells (**g-i**). Cells were transfected with siRNAs targeting IFITM proteins and scrambled siRNA (NC). IFN-α2b was used to stimulate the expression of IFITM proteins. The mRNA expression levels were detected using qPCR and normalized to that of NC at the untreated condition (-IFN-α). **d** and **j**, endogenous IFITM1,2,3 proteins expression in HuMEC (**d**) and HEK293T cells (**j**) upon siRNA knockdown analyzed by Western Blot. IFITMs were detected by anti-IFITM antibodies, and GAPDH was a loading control. The entry of NiV/VSV pp and HeV/VSV pp to HuMEC (**e** and **f**) and HEK293T cells (**k** and **l**). NiV and HeV glycoproteins were pseudotyped to VSV particles in which the VSV-G gene was replaced with the Renilla luciferase gene. Virus entry was measured by luminescence intensity and normalized to that of scrambled siRNA at the untreated condition (NC, −IFN-α). Virus entry levels in siRNA-transfected cells were compared with those transfected with scrambled siRNA (NC) under respective −IFN-α and +IFN-α conditions. Virus entry in IFN-α–treated cells (+IFN-α) was further compared with that in resting cells (−IFN-α) and labeled with a bracket. Bars represent means ± SEM. Results from at least 3 independent experiments are shown. *p* values were obtained using one-way analysis of variance (ANOVA) with post hoc correction (nonsignificant [ns], *p* > 0.05; *, *p* ≤ 0.05; **, *p* ≤ 0.01; ***, *p* ≤ 0.001; ****, *p* ≤ 0.0001).

We next measured the entry of NiV- and HeV-glycoproteins pseudotyped vesicular stomatitis virus particles (NiV/VSV pp and HeV/VSV pp, respectively) into untreated or IFN-α–stimulated HuMEC and HEK293T cells (Fig. 1 e,f,k and l). To validate the VSV pseudovirus system, we pseudotyped HA and NA of IAV (Thailand KAN-1/2004 H5N1 strain) and env of MLV (10A1 strain) to VSV, and confirmed that the entry of IAV/VSV pp into HEK293T cells is inhibited upon IFITM2 and IFITM3 overexpression, but not IFITM1 (Fig. S2a), whereas MLV/VSV pp entry is unaffected by the overexpression of any IFITM protein (Fig. S2b), consistent with previous reports (23, 36). IFN-α treatment significantly inhibited the entry of both NiV/VSV pp and HeV/VSV pp in HuMEC cells (NC, Fig. 1e and f) and HEK293T cells (NC, Fig. 1k and l). In HuMEC, IFITM1-knockdown resulted in a significant enhancement of both NiV/VSV pp and HeV/VSV pp entry in the untreated and IFN-α stimulated conditions (Fig. 1e and f), while knockdown of IFITM2 and IFITM3 led to slight to no increase (Fig. 1e and f). In HEK293T cells, knockdown of IFITM1 and IFITM2 significantly increased the entry of both NiV/VSV pp and HeV/VSV pp in the untreated and IFN-α stimulated conditions (Fig. 1k and 1l), while only a moderate increase in the infection of NiV/VSV pp and HeV/VSV pp was observed with IFITM3 knockdown. Our results suggest that endogenous IFITM1 is the most potent in restricting HeV and NiV entry among the three IFITM proteins, which is consistent with previous findings that IFITM1 inhibits infection of paramyxoviruses that enter at the plasma membrane (6).

**Fig. S1.**
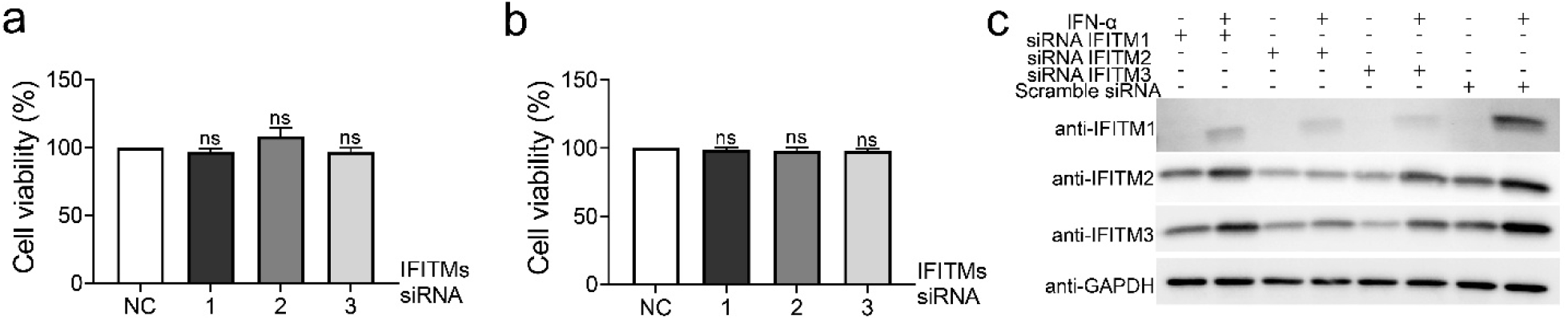
**a** and **b**, the viability of HEK293T cells (**a**) and HuVEC (**b**) treated with IFITMs siRNAs was detected using a CCK8 kit. Bars represent means ± SEM. Results from at least three independent experiments are shown. *p* values were obtained using one-way analysis of variance (ANOVA) with post hoc correction (nonsignificant [ns], *p* > 0.05). **c**, Cell lysates were harvested from HEK293T cells transfected with siRNA targeting IFITM1, 2, 3, or a scramble. Endogenous IFITM1, 2, and 3 were analyzed using Western Blot. IFITMs were detected using anti-IFITMs antibodies and GAPDH served as a loading control.

**Fig. S2.**
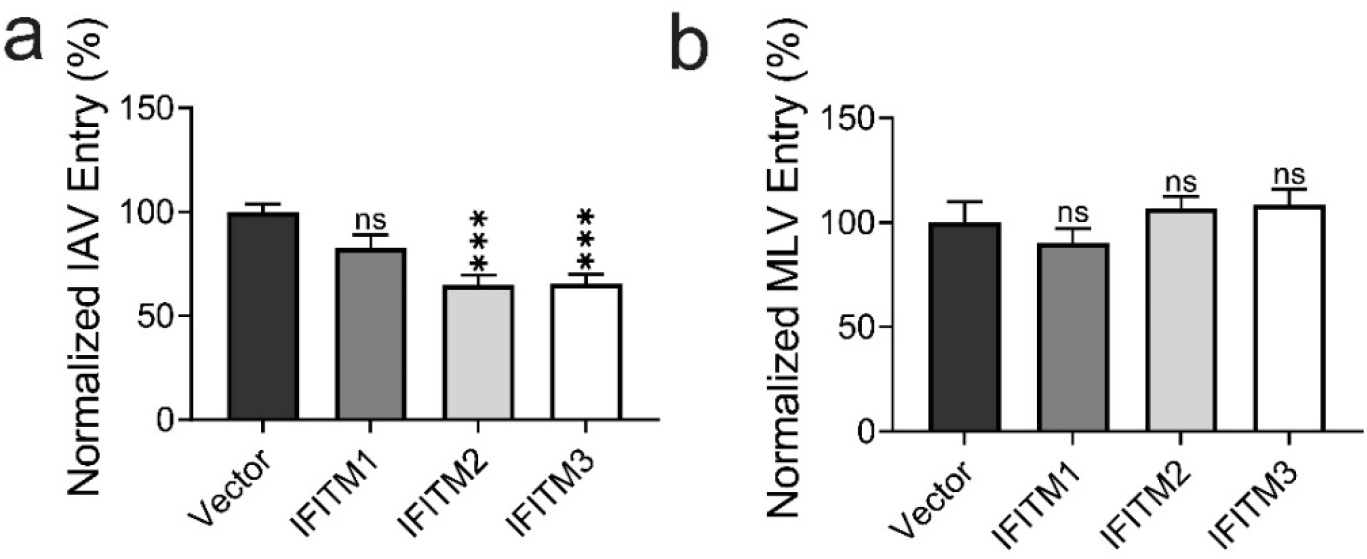
The entry of IAV/VSV pp (**a**) and MLV/VSV pp (**b**) in HEK293T cells stably expressing IFITM proteins. HEK293T cells transduced with an empty vector retrovirus were used as a control (Vector). Virus entry was measured by luminescence intensity and normalized to that of Vector. Bars represent means ± SEM. Results from at least three independent experiments are shown. *p* values were obtained using one-way analysis of variance (ANOVA) with post hoc correction. (nonsignificant [ns], *p* > 0.05; ***, *p* ≤ 0.001).

Next, we investigated whether endogenous IFITM proteins also inhibit cell-cell fusion caused by NiV and HeV glycoproteins in endothelial and epithelial cells. Fusion proteins of paramyxoviruses execute membrane fusion at neutral pH, thus can cause significant cell-cell fusion (syncytia), which is a pathological hallmark of infection and enables efficient cell-to-cell spread of the virus (37, 38). Additionally, multinucleated syncytia caused by HNV infection lead to vasculitis in the central nervous system, which gives rise to lethal encephalitis (33, 34). We used a heterologous fusion assay to examine the role of endogenous IFITMs in HNV-induced cell-cell fusion. We chose PK13 cells as effector cells because these cells lack endogenous ephrinB2 receptors and thus do not self-fuse upon expression of HNV glycoproteins (30, 39). Effector cells transfected with HeV or NiV glycoproteins were lifted off and overlaid on target cells transfected with IFITMs siRNAs either being untreated or IFN-α-stimulated. In human umbilical vein endothelial cells (HuVECs), siRNA-mediated knockdown of IFITM1, IFITM2, and IFITM3 significantly reduced both IFITM mRNA levels (Fig. S3a-c) and protein levels (Fig. S3d). In the untreated HuVECs cells, knockdown of IFITM1 and IFITM2 enhanced NiV-induced cell-cell fusion (Fig. 2a), while knockdown of IFITM1, IFITM2, and IFITM3 proteins promoted HeV-induced cell-cell fusion (Fig 2b). Intriguingly, neither NiV-nor HeV-induced cell-cell fusion was inhibited upon IFN-α stimulation (Fig. 2a and 2b), despite the elevated mRNA and protein expression of IFITMs (Fig. S3). It is possible that IFN-α either up- or down-regulates other factors that counteract IFITM activity, or alters the post-translational modification of IFITMs by enzymes such as ZDHHCs (40) and ABHD16A (41). In both untreated and IFN-α-stimulated HEK293T cells, knockdown of IFITM1, 2, and 3 promoted NiV-induced cell-cell fusion (Fig. 2c), while only knockdown of IFITM1 and IFITM2 led to an enhancement of HeV-induced cell-cell fusion (Fig. 2d). Therefore, endogenous IFITM1 and IFITM2 appear to play a major role in restricting NiV- and HeV-induced cell–cell fusion, whereas IFITM3 contributes to a lesser extent. We confirmed that the siRNA knockdown of IFITM proteins does not affect the levels of ephrinB2 (Fig. S4a). This finding is consistent with the role of endogenous IFITM proteins in restricting HNV virus entry (Fig. 1g-l).

**Fig. 2.**
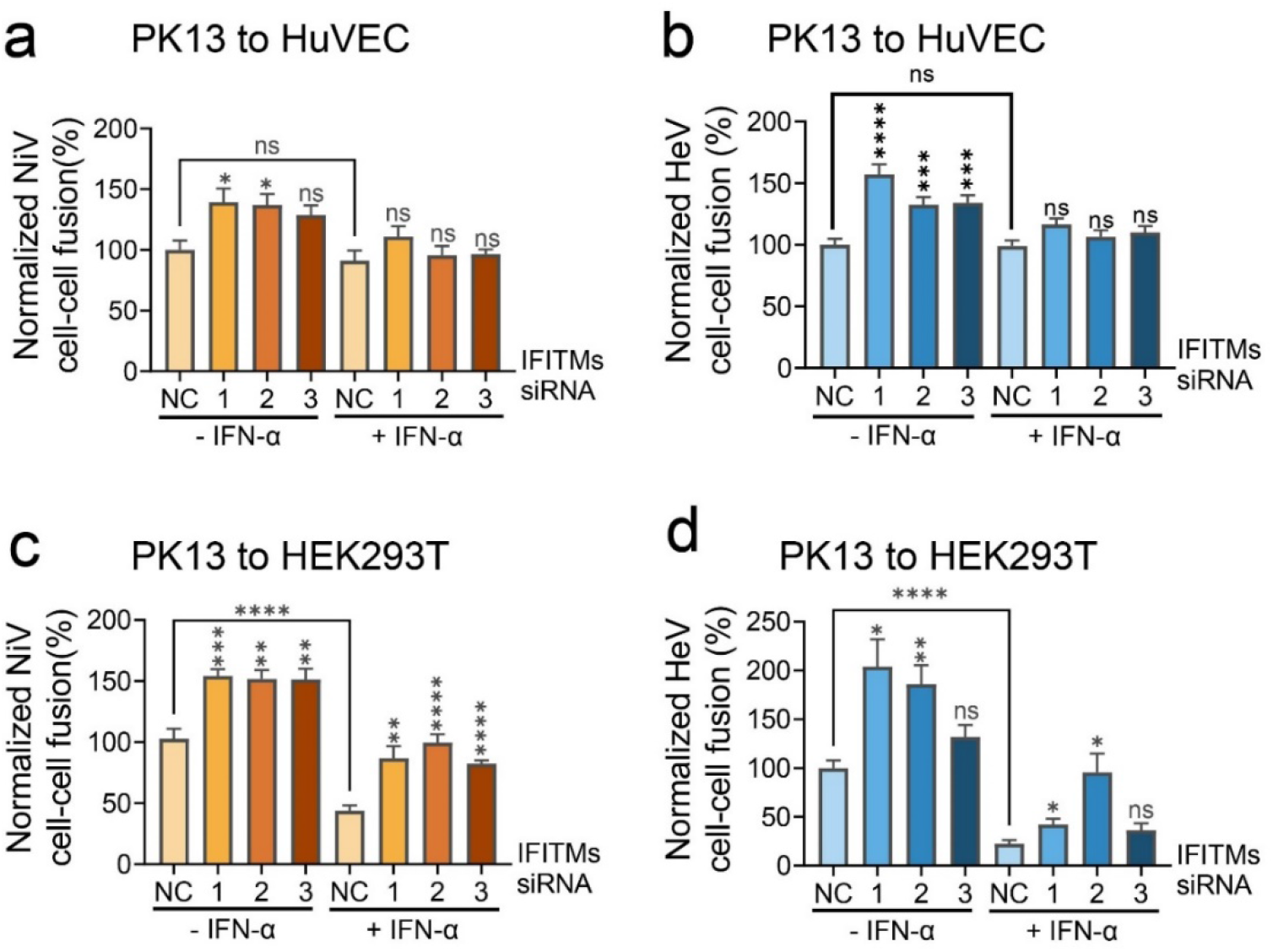
The role of endogenous IFITMs in NiV and HeV-induced cell-cell fusion. PK13 cells expressing NiV-F and G (**a** and **c**) or HeV-F and G (**b** and **d**) were used as effector cells. Target HuVEC (**a** and **b**) and HEK293T (**c** and **d**) cells were transfected with siRNA targeting IFITMs. After-transfection, target cells were treated overnight with IFN-α2b to induce the expression of IFITM proteins. Effector cells were lifted and overlaid on target cells at 24 hrs post-transfection with HNV-F and G. After co-culture, cells were fixed, and syncytia were quantified by counting the number of nuclei in the syncytium per 10 × field (5 fields are counted per group) under a Nikon TE2000U microscope. All data were normalized to that of scrambled siRNA at untreated condition (NC, −IFN-α). Cell-cell fusion levels in siRNA-transfected cells were compared with those transfected with scrambled siRNA (NC) under respective −IFN-α and +IFN-α conditions. Cell-cell fusion in IFN-α–treated cells (+IFN-α) was further compared with that in the untreated condition (−IFN-α). Bars represent means ± SEM. Results from at least three independent experiments are shown. *p* values were obtained using one-way analysis of variance (ANOVA) with post hoc correction. (nonsignificant [ns], *p* > 0.05; *, *p* ≤ 0.05; **, *p* ≤ 0.01; ***, *p* ≤ 0.001; ****, *p* ≤ 0.0001).

**Fig. S3.**
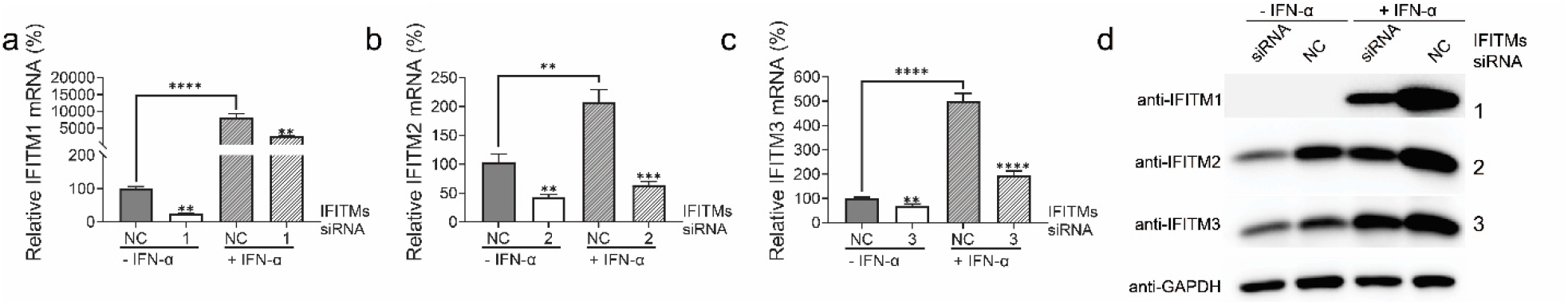
IFITM knockdown in HuVEC. **a-c**, the endogenous IFITMs mRNA levels in HuVEC transfected with separate siRNA targeting IFITM proteins and scrambled siRNA (NC). IFN-α2b was used to stimulate the expression of IFITM proteins. The mRNA expression levels were detected using qPCR and normalized to that of NC untreated by IFN-α (-IFN-α). *p* values were obtained using one-way analysis of variance (ANOVA) with post hoc correction (**, *p* ≤ 0.01; ***, *p* ≤ 0.001; ****, *p* ≤ 0.0001). Bars represent means ± SEM. Results from at least 3 independent experiments are shown. **d**, endogenous IFITM proteins expression in HuVEC upon siRNA knockdown was analyzed by Western Blot. IFITMs were detected by anti-IFITM antibody, and GAPDH served as a loading control.

### IFITMs overexpression affects HNV glycoprotein-induced membrane fusion in a cell type-dependent manner

To further determine the role of IFITMs in HNV-mediated membrane fusion, we established HeLa and HEK293T cell lines that stably express FLAG-tagged IFITMs. The expression of FLAG-IFITMs in HEK293T (Fig. 3a) and HeLa (Fig. 3b) cells was confirmed by western blots. The subcellular localization of IFITMs in HEK293T (Fig. 3c) and HeLa (Fig. 3d) cells agrees with previous results (4), with IFITM1 primarily on the plasma membrane, IFITM2 in the endosomes, and IFITM3 on both the cell surface and endosomes. Overexpression of IFITM1 in HEK293T cells reduced the entry of NiV/VSV pp and HeV/VSV pp by about 50%, while overexpression of IFITM2 or IFITM3 led to a lesser but significant reduction of VSV/NiV pp and VSV/HeV pp entry (Fig. 3e and 3f). Similarly, a significant inhibition of NiV-(Fig. 3g) and HeV- (Fig. 3h) glycoproteins-induced cell-cell fusion by IFITM proteins was observed in HEK293T cells, with a ∼80% inhibition of IFITM1 against NiV glycoproteins-induced cell-cell fusion. These results suggest that IFITM1 is in general the most potent in restricting NiV and HeV-mediated virus entry and cell-cell fusion in HEK293T cells, consisting with the observation for endogenous IFITMs (Fig. 1k,1l, 2c and 2d). In HeLa cells, we noticed that overexpression of IFITM1 and IFITM3 inhibits the entry of VSV/NiV pp (Fig. 3i) and VSV/HeV pp (Fig. 3j), while overexpression of IFITM2 enhances the entry of VSV/NiV pp (Fig. 3i) and VSV/HeV pp (Fig. 3j). It is plausible that the overexpressed IFITM2 in HeLa cells is more localized to endosomes, thus may facilitate the entry of HNV/VSV pp via macropinocytosis by directly interacting with the viral glycoprotein in endosomes (12, 42). IFITM proteins significantly inhibited cell-cell fusion induced by NiV (Fig. 3k) and HeV glycoproteins (Fig. 3l) in HeLa cells. We confirmed that overexpression of IFITMs did not change ephrinB2 expression (Fig. S4b). Taken together, overexpression of IFITM1 and IFITM3 restricts NiV and HeV-induced virus entry and cell-cell fusion in HEK293T and HeLa cells.

**Fig. 3.**
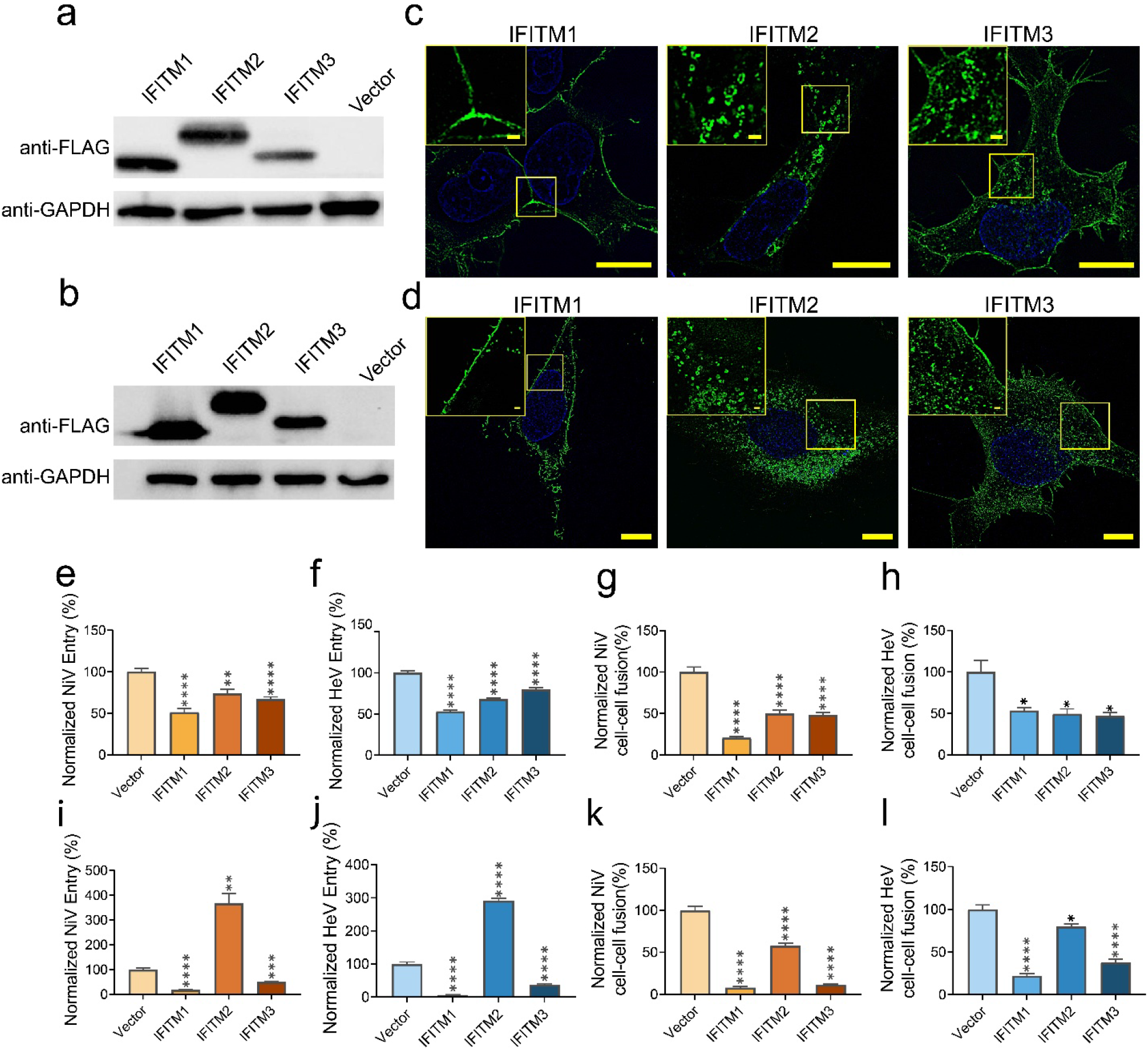
IFITMs overexpression affects NiV and HeV entry and cell-cell fusion differently. The protein expression levels of HEK293T cells (**a**) and HeLa cells (**b**) stably express IFITMs. HEK293T or HeLa cells were transduced with retroviruses encoding FLAG-tagged IFITM proteins in the pQCXIP vector or with an empty vector control, followed by puromycin selection. Cells were analyzed by Western Blot. IFITMs were detected using an anti-FLAG antibody, and GAPDH served as a loading control. Fluorescence microscopy images of HEK293T (**c**) and HeLa (**d**) cells overexpressing IFITM proteins. Cells were imaged using fluorescence microscopy. IFITMs were stained with a mouse anti-FLAG primary antibody and a donkey anti-mouse Alexa Fluor 488 secondary antibody. The cell nuclei were stained with DAPI. Scale bars: 10 µm. The boxed regions are enlarged. Scale bar: 1 μm. The entry of NiV/VSV pp and HeV/VSV pp to HEK293T cells (**e** and **f**) and HeLa cells (**i** and **j**) stably expressing IFITMs, respectively. HNV/VSV pp were made as described above, and virus entry was measured by luminescence intensity and normalized to that of Vector. Cell-cell fusion induced by NiV and HeV glycoproteins in HEK293T (**g** and **h**) and HeLa cells (**k** and **l**) stably expressing IFITMs. NiV-F and G or HeV-F and G were co-transfected into HEK293T-IFITMs and HeLa-IFITMs cells. At 18-24 hrs post-transfection, cells were fixed, and syncytia were quantified by counting the number of nuclei in the syncytium per 10 × field (5 fields are counted per group). Cell-cell fusion levels were normalized to those of Vector. Bars represent means ± SEM. Results from at least three independent experiments are shown. *p* values were obtained using one-way analysis of variance (ANOVA) with post hoc correction (*p* > 0.05; *, *p* ≤ 0.05; **, *p* ≤ 0.01; ***, *p* ≤ 0.001; ****, *p* ≤ 0.0001).

**Fig. S4.**
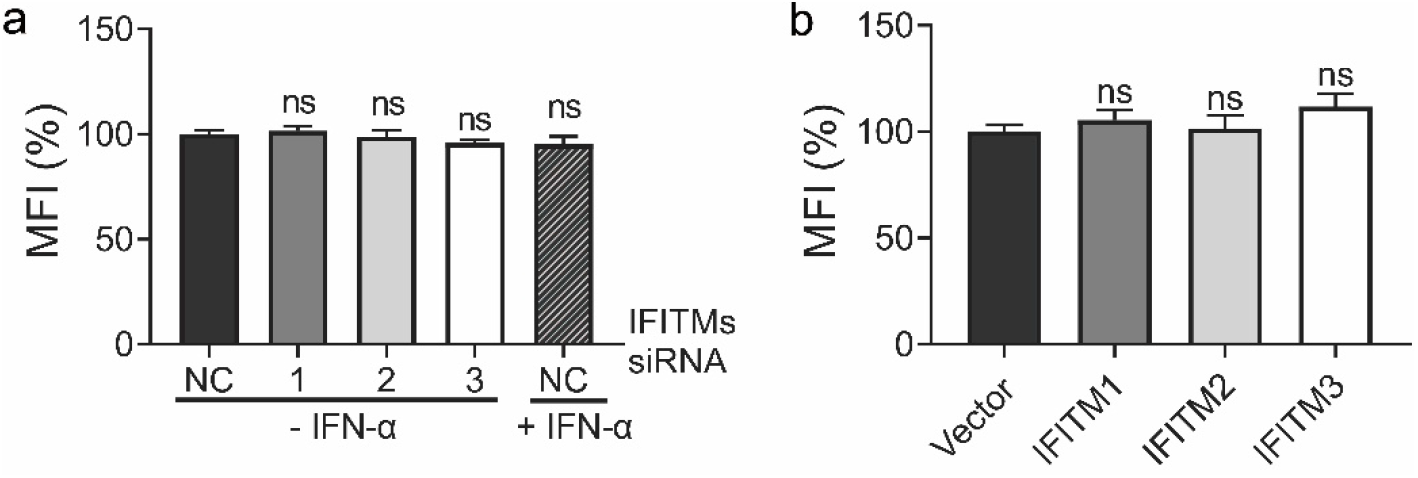
The knockdown and overexpression of IFITM proteins do not affect the cell surface levels of ephrinB2 in HEK293T cells. **a**, the expression of endogenous ephrinB2 in IFITMs-knockdown HEK293T cells. HEK293T cells were transfected with siRNA targeting IFITM proteins or treated with IFN-α2b to stimulate the expression of IFITM proteins. EphrinB2 on the cell surface was detected by a recombinant human EphB3-Fc chimera and AlexaFluor 488-conjugated donkey anti-human antibody. The mean fluorescence intensity (MFI) of AlexaFluor 488 was determined by flow cytometry and normalized to that of Vector. **b**, the expression of endogenous ephrinB2 in HEK293T cells stably expressing IFITM proteins. EphrinB2 was detected as above. Results from at least three independent experiments are shown. Bars represent means ± SEM. *p* values were obtained using one-way analysis of variance (ANOVA) with post hoc correction (nonsignificant [ns], *p* > 0.05).

### IFITM1 forms nanoclusters on the cell surface

Next, we investigated how IFITM1 inhibits HNV-induced membrane fusion. IFITM proteins broadly inhibit viral entry by modulating membrane properties. IFITM3 has been shown to cluster on endosomal membranes (43, 44), and IFITM1 on plasma membranes by super-resolution microscopy (45). Furthermore, oligomerization of IFITM3, governed by a 91-GxxxG-95 motif, is critical for its antiviral activity by increasing membrane stiffness (13). Compared with IFITM3, IFITM1 lacks the N-terminal 21 amino acids that include an endocytic YxxΦ motif (4). As a result, IFITM1 predominantly localizes to the plasma membrane, and inhibits HNV that enters at the cell surface. We first probed the nano-organization of endogenous IFITM1 on the plasma membrane in HEK293T cells. HEK293T cells were selected because 1) the basal level of endogenous IFITM1 is low and not detectable by western blot (Fig. 1j), thus providing a background control to resolve IFN-induced IFITM1 molecular reorganization; 2) the overexpressed and endogenous IFITM1 showed consistent inhibitory effect on HNV-induced membrane fusion (Fig. 1 and 3). To investigate the nano-organization of IFITM1 on the plasma membrane, we performed single-molecule localization microscopy (SMLM) on the ventral membrane of HEK293T cells under the untreated and IFN-stimulated conditions. The custom-built SMLM system achieves < 10 nm localization precision in the lateral direction (46, 47). Upon IFN stimulation, the IFITM1 clusters become more prominent (Fig. 4b) than that in the untreated cells (Fig. 4a). Cluster identification was performed using DBSCAN (Density-Based Spatial Clustering of Applications with Noise), which links the closely situated localizations in a propagative manner (cluster maps, Fig. 4a and b) (46, 48). We show that IFITM1 molecules are more likely to distribute into clusters upon IFN treatment, demonstrated by a higher Hopkins’ index (Fig. 4c) and percentage of localizations in clusters (Fig. 4d). IFITM1 clusters are larger (Fig. 4e) but with similar molecule density (Fig. 4f). We also noticed that more IFITM1 clusters are formed upon IFN stimulation (Fig. 4g). These results reveal the clustering pattern of IFITM1 on a nanoscale, and its clustering is enhanced upon IFN stimulation.

**Fig. 4.**
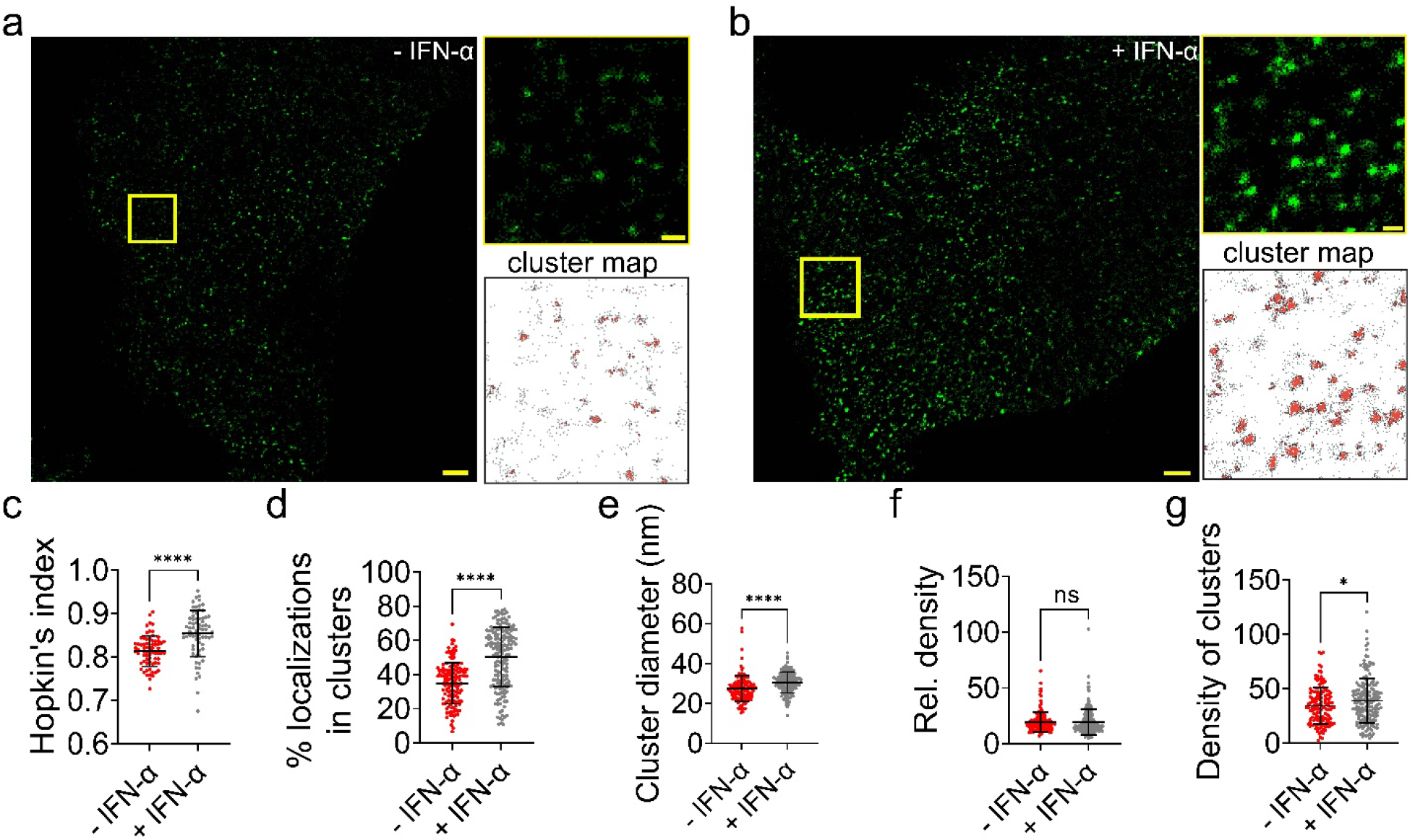
Endogenous IFITM1 forms clusters on HEK293T cells. IFN-α2b- and mock-treated HEK293T cells were fixed and permeabilized. IFITM1 was probed using a mouse anti-IFITM1 monoclonal antibody and an Alexa Fluor 647-conjugated goat anti-mouse antibody. **a** and **b**, x-y cross-section (600 nm thick in z) of the SMLM images of endogenous IFITM1 at the ventral membrane of control (**a**, - IFN-α) and IFN-α2b-treated (**b**, + IFN-α) HEK293T cells. Scale bar: 1 μm. The boxed regions are enlarged. Scale bar: 200 nm. The cluster maps are shown. Cluster contours are in gray. **c** to **g**, the Hopkin’s index (A high value indicates clustering tendency.) (**c**), percentage of localizations in clusters (**d**), cluster diameter (**e**), relative density (the ratio of the localization density within clusters to that of the total region of interest (ROI)) (**f**), and the density of clusters (**g**) are shown in dot plots. Sample size n = 141 (-IFN-α) and 173 (+ IFN-α) from 7 cells per group. Bars represent means ± SD. *p* value was obtained using Welch’s *t*-test. (nonsignificant [ns], *p* > 0.05; *, *p* ≤ 0.05; ****, *p* ≤ 0.0001).

### IFITM1 sequesters ephrinB2 in raft-like membrane domains

IFITM3 clustering is a determinant of its antiviral activity and has been associated with increased membrane order (13). Additional studies show that both IFITM1 and IFITM3 either target raft-like membrane domains by S-palmitoylation (16, 49) or sort membrane lipids by directly interacting with cholesterol (15, 26). Cholesterol is enriched in nanoscale membrane domains that are thought to exhibit raft-like, liquid-ordered character in the plasma membrane (50, 51). We hypothesize that IFITM1 clusters restrict HNV entry by sequestering ephrinB2 within rigid membrane nanodomains, thereby increasing the energy barrier in virus-cell membrane fusion. To test this hypothesis, we started by probing the co-occurrence of IFITM1 clusters and liquid-ordered membrane domains. Cholera toxin B (CtxB) is widely used to label GM1 (monosialotetrahexosylgangliosides)-enriched, cholesterol-dependent membrane nanodomains that are often referred as lipid rafts (52). To determine the spatial relationship of IFITM1 to GM1 lipid domains, we performed two-color SMLM imaging. The fixation process, employing 4% paraformaldehyde and 0.2% glutaraldehyde, was applied before and after CtxB staining, ensuring that protein movement and CtxB-binding-induced lipid domain collapse were minimized (52, 53). The results showed that GM1 clusters appear larger and more crowded than IFITM1 clusters (Fig. 5a). This may partly reflect the different abundance of GM1 domains and IFITM1 assemblies on the plasma membrane, but it may also result from differences in the brightness and photo-switching property of fluorophores used to label GM1 and IFITM1 (54). Accordingly, the two-color SMLM data mainly inform the relative spatial distribution of GM1 and IFITM1, whereas cluster properties from two channels should not be interpreted as the absolute organization of either molecules or compared between channels. We also noticed that many GM1 clusters are closely spaced and appear to form larger GM1-enriched cluster islands (Fig. 5a, degree of colocalization map). Within these structures, IFITM1-associated GM1 localizations are concentrated at the periphery, instead of spreading throughout (Fig. 5a, degree of colocalization map, black arrows). Meanwhile, the much smaller IFITM1 clusters were often either entirely colocalized with GM1 or separated from it. We also noticed that a number of IFITM1 localizations do not colocalize with GM1 (gray localizations in cluster map, Fig. 5a), and these localizations either do not form clusters or partition into much smaller clusters (Fig. 5a). This pattern implies the possibility that IFITM1 preferentially associates with the edge of GM1 domains. However, our data cannot distinguish whether a GM1-enriched domain corresponds to a GM1 cluster or a larger cluster island. Consistently, our analysis shows that both IFITM1 clusters and GM1 clusters are enlarged when associated with one another, suggesting a non-random spatial co-occurrence between IFITM1 clusters and GM1 clusters (Fig. 5a-c). The relative density of the GM1 localizations is elevated in IFITM1-associated clusters, suggesting that IFITM1 may either recruit GM1 laterally or target GM1 concentrated domains, consisting with the notion that IFITM1 is both *S*-palmitoylated and can directly interact with cholesterol (Fig. 5d) (15, 16). The GM1-associated IFITM1 clusters are not more densely packed than the free IFITM1 clusters (Fig. 5e), indicating that the larger GM1-associated IFITM1 clusters may result from a collapse of nearby small IFITM1 clusters, instead of re-organization of molecules inside those clusters. Nonetheless, this finding suggests that IFITM1 clusters preferentially associate with GM1-enriched membrane domains, which may represent more ordered, rigid membrane regions.

**Fig. 5.**
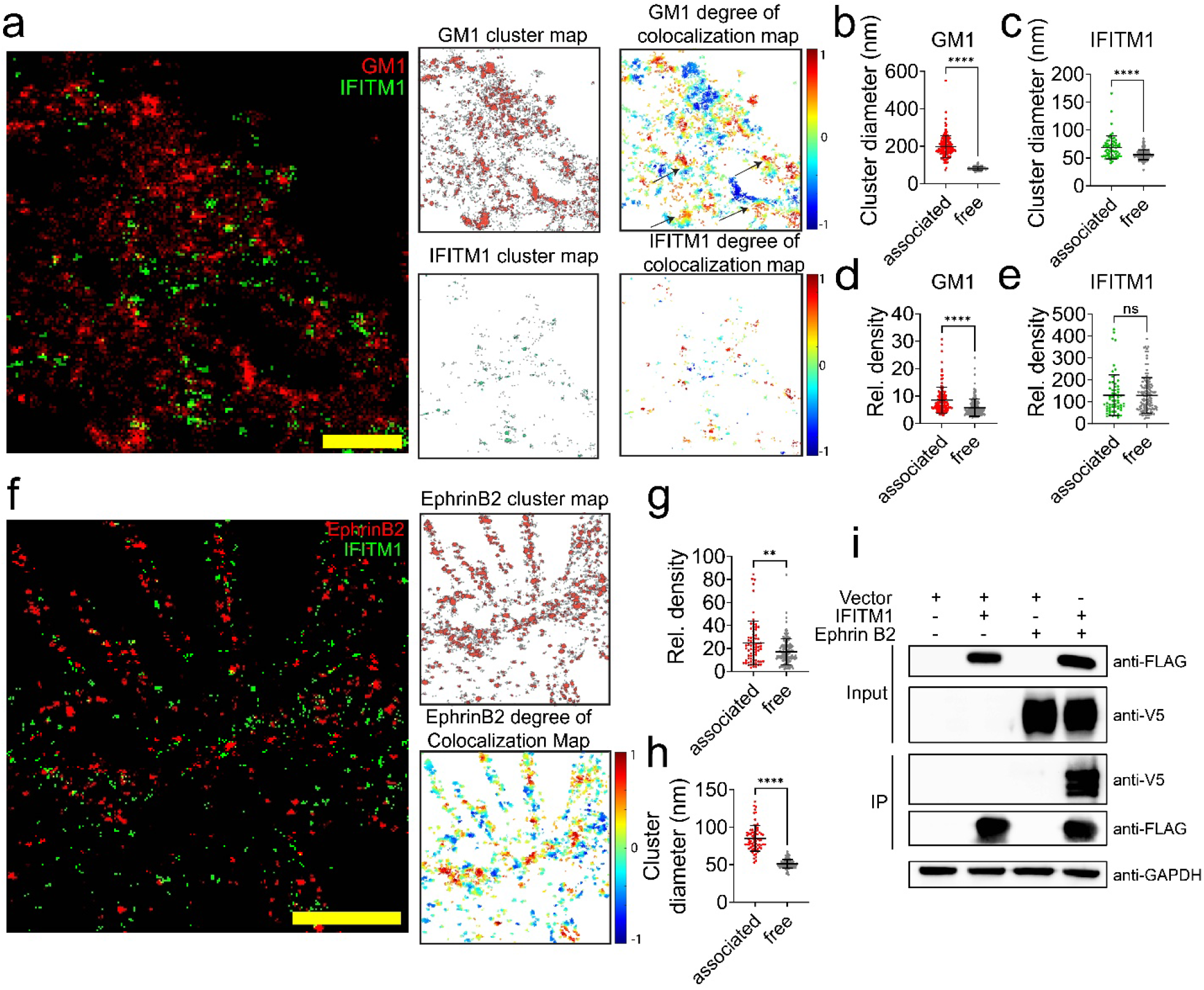
IFITM1 sequesters ephrinB2 in GM1 membrane domains. **a** to **e**, HEK293T cells stably expressing FLAG-tagged IFITM1 were fixed and stained with cholera toxin subunit B (ctxB) conjugated to 647 (detect GM1), followed by a second fixation to immobilize the bound ctxB. Cells were then permeabilized and stained with mouse anti-Flag antibody and cy3B-conjugated donkey anti-mouse antibody (detect IFITM1). **a**, a x-y cross-section (600 nm thick in z) of an SMLM image of IFITM1 (green) and GM1 (red) on the ventral plasma membrane, the cluster maps and degree of colocalization maps of GM1 and IFITM1 localizations are shown. In the degree of colocalization maps, each localization is color-coded by its degree of colocalization value according to the scale on the right (see *Materials and Methods*.). **b** to **e**, the cluster diameter and relative density of GM1 clusters (**b** and **d**) and IFITM1 clusters (**c** and **e**) are shown in dot plots. See “*Materials and Methods*” for the classification of *associated* and *free* clusters. For GM1 clusters, n = 138 (associated) and 169 (free) from 8 cells. For IFITM1 clusters, n = 62 (associated) and 125 (free) from 8 cells. **f** to **h**, HEK293T cells stably expressing FLAG-tagged IFITM1 were transfected by plasmids coding for V5-tagged ephrinB2. At 24 hrs post-transfection, cells were permeabilized and stained with mouse anti-FLAG antibody and cy3B-conjugated donkey anti-mouse antibody (detect IFITM1), and goat anti-ephrinB2 antibody and AlexaFluor 647-conjugated donkey anti-goat antibody (detect ephrinB2). **f**, A x-y cross-section (600 nm thick in z) of an SMLM image of IFITM1 (green) and ephrin B2 (red) on the ventral plasma membrane, the cluster map, and the degree of colocalization map of ephrinB2 localizations. **g and h**, the relative density (**g**) and cluster diameter (**h**) of the free and IFITM1-associated ephrinB2 clusters are shown in dot plots. Sample size n = 70 (associated) and 135 (free) from 8 cells per group. Bars represent means ± SD. *p-* value was obtained using Welch’s *t*-test. (nonsignificant [ns], *p* > 0.05; **, *p* ≤ 0.01; ****, *p* ≤ 0.0001). **i**, ephrinB2 is co-immunoprecipitated with IFITM1. HEK293T cells were transfected with empty vector and plasmids coding for IFITM1-FLAG and ephrinB2-V5. IFITM1 was immunoprecipitated using anti-FLAG magnetic beads. Input and IP samples were analyzed by SDS-PAGE and immunoblotted with mouse anti-FLAG and rabbit anti-V5 antibodies.

Next, we investigated whether IFITM1 sequesters ephrinB2 into the GM1 domains. SMLM images of ephrinB2 on HEK293T plasma membrane show that ephrinB2 molecules form clusters, with most selected regions showing a Hopkin’s index > 0.9 (Fig. S5a and S5b) and a cluster diameter of 44-60 nm (Fig. S5c). We performed dual-color SMLM on IFITM1 and ephrinB2 on the plasma membrane of 293T cells to visualize the spatial distribution and organization of ephrinB2 relative to IFITM1. Images show that some ephrinB2 clusters overlap with IFITM1 (Fig. 5f). The co-clustering between ephrinB2 and IFITM1, marked by orange-red dots in the degree of colocalization map, was mainly observed at the cell periphery and filopodia, where IFITM1 is predominantly expressed (Fig. 5f). Our results show that the relative density of ephrinB2 localizations is higher in IFITM1-associated ephrinB2 clusters than in free clusters (Fig. 5g), and IFITM1-associated ephrinB2 clusters are larger than free clusters (Fig. 5h). Consistently, the co-IP assay confirms interactions between ephrinB2 and IFITM1 (Fig. 5i). These data suggest that IFITM1 reorganizes ephrinB2 clusters into larger and more densely packed assemblies without directly enhancing the bulk ephrinB2 expression (Fig. S4b). Given the co-occurrence of IFITM1 with GM1 domains, we speculate that virus-cell membrane fusion triggered through receptors in these regions may be disfavored, potentially because of the enhanced rigidity of the receptor-associated membrane domain.

**Fig. S5.**
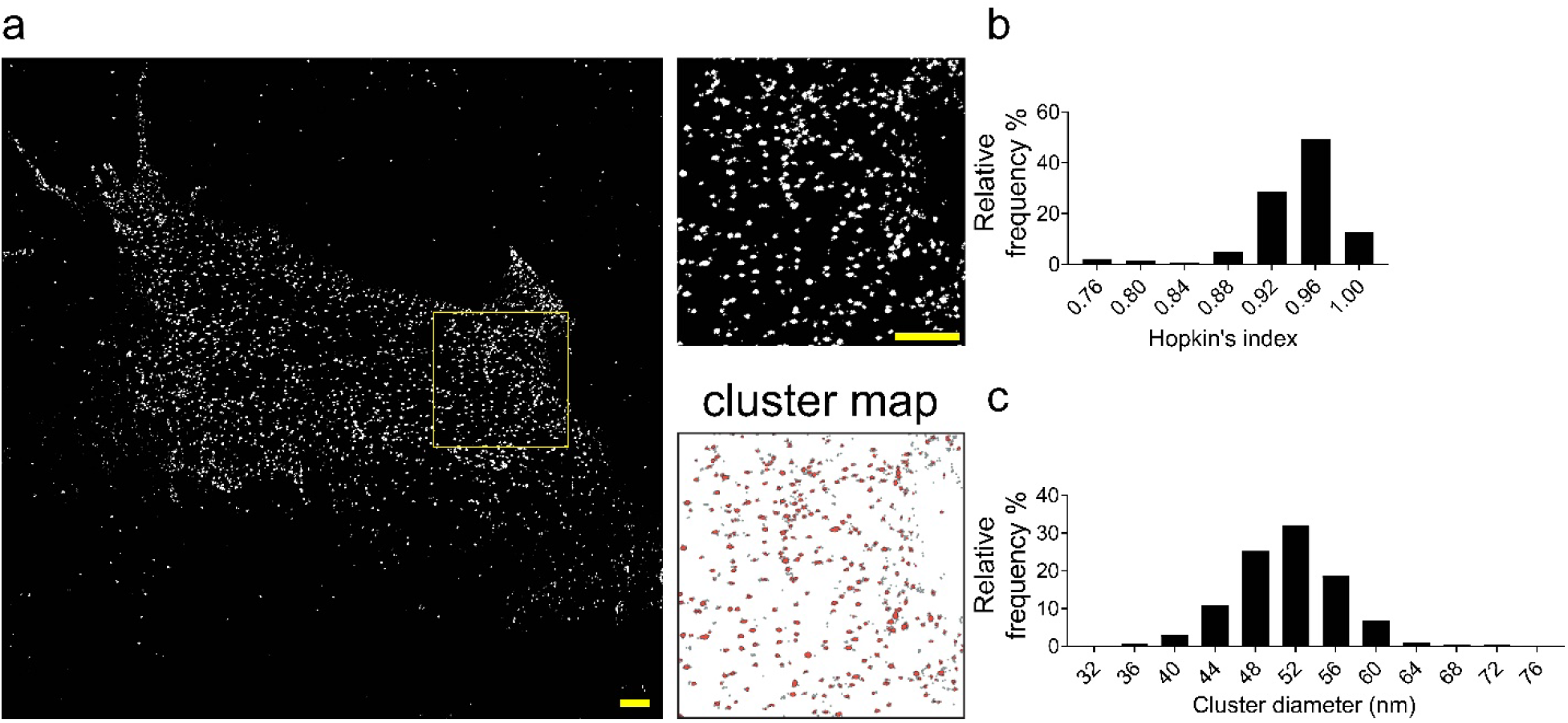
EphrinB2 molecules cluster on HEK293T cells. HEK293T cells over-expressing V5-tagged ephrinB2 were fixed and stained by using goat anti-ephrinB2 antibody and AlexaFluor 647-conjugated donkey anti-goat antibody. **a**, a *x-y* cross-section (Δz = 600 nm) of SMLM images of ephrin B2 on the ventral membrane of 293T cells. Scale bar: 1 μm. The yellow boxed region is enlarged to show the detailed distribution pattern. Scale bar: 1 μm. A cluster map of ephrinB2 localizations is shown. **b**, Relative frequency distribution of the Hopkin’s index for ephrinB2 localizations. Sample size n = 140 from 14 cells. **c**, Relative frequency distribution of the ephrinB2 cluster diameter. Sample size n = 327 from 14 cells.

### IFITM1 restricts ephrinB2 lateral diffusion on the plasma membrane

We next asked whether the IFITM1-associated GM1 domains confine ephrinB2 mobility. Because ephrinB2 is a transmembrane protein, its lateral diffusion in the plasma membrane is expected to depend on the local membrane environment. We therefore reasoned that IFITM1-mediated partitioning of ephrinB2 into GM1 domains, presumably more-ordered, less fluid, could reduce its movement on the cell surface (55, 56). Because previous studies have primarily focused on IFITM3 for membrane rigidification, we used IFITM3 as a control. In our system, IFITM3 inhibits both HNV entry and cell-cell fusion in HEK293T cells, albeit to a lesser extent than IFITM1 (Fig. 1 and 3). We performed live-cell imaging and single-particle tracking to analyze the ephrinB2 mobility on the plasma membrane. EphrinB2 was sparsely labeled with mStaygold at its C-terminus and transfected into HEK293T cells stably expressing IFITMs (293T-IFITMs). The sparse labeling is achieved by a translational readthrough sequence inserted between the open reading frames for ephrinB2 and mStaygold, allowing only 0.4% of ephrinB2 molecules to be labeled by mStaygold (57). This allows us to track individual ephrinB2 clusters that would otherwise appear as a continuous fluorescent signal delineating the membrane. Additionally, the superior photostability and brightness of mStaygold allows for long-term, high frame-rate tracking (58). Interestingly, ephrinB2 moves less freely in 293T-IFITM1 and 293T-IFITM3 cells than control, as evidenced by more confined tracks (Fig. 6a and b). We noticed the heterogeneity in the motion states of individual molecules (Fig. 6b, vector). To precisely analyze the motions, we applied a transient motion analysis approach DC-MSS (divide-and-conquer moment scaling spectrum), which allows for the identification of three distinct mobility states (immobile, confined, and free) along each track (Fig. 6c). Our analysis results show that ephrinB2 molecules are much more confined in 293T-IFITM1 cells than in control cells, marked by an enhanced fraction of time in confined state at the expense of the free state (Fig.6d). Interestingly, IFITM3 shows a shift in the fraction of time from an immobile state to a confined state, while the time spent in free motion remains similar to the control cells (Fig. 6d). This indicates that ephrinB2 predominantly exhibits confined diffusion when IFITM1 is overexpressed, whereas the distribution of ephrinB2 motion states was affected to a lesser extent by IFITM3. Under confined and immobile conditions, ephrinB2 molecules move within a smaller area in 293T-IFITM1 and 293T-IFITM3 cells than control (Fig. 6e). Similarly, ephrinB2 molecules show lower diffusion coefficients in 293T-IFITM1 and 293T-IFITM3 cells than control (Fig. 6f). This data strongly suggests that the spatial movement of ephrinB2 is reduced upon IFITM1 and IFITM3 overexpression. To further test whether the reduced ephrinB2 motion could be attributed to increased rigidity of the membrane domains surrounding ephrinB2, we thought to enhance the bulk membrane fluidity using amphotericin B (ampho B). Previous study shows that ampho B counteracts IFITM3-mediated restriction on IAV by increasing membrane fluidity (13, 24). Therefore, it is plausible that ampho B treatment may increase ephrinB2 lateral diffusion and thus counteract IFITM1’s restriction on NiV entry. In our system, ampho B treatment rescues ephrinB2 restriction by IFITM3, but not IFITM1, demonstrated by tracks (Fig. 6g and h) and diffusion coefficients (Fig. 6i). Consistently, ampho B treatment counteracted IFITM3-mediated NiV restriction, but was not effective on IFITM1-mediated NiV restriction (Fig. 6j). It is worth noting that ampho B counteracts IFITM3-mediated IAV restriction, but not IFITM1; and IAV mainly enters via endocytosis (24). These data suggest that 1) the confined movement of ephrinB2 in 293T-IFITM3 cells may result from decreased membrane fluidity, as it can be rescued by ampho B treatment; 2) IFITM1-mediated restriction of ephrinB2 lateral mobility is not solely attributed to enhanced membrane rigidity, or that such rigidification is insensitive to amphotericin B treatment. This is not surprising because the regulation of protein diffusion by raft partitioning or cholesterol is context-dependent rather than universal (55, 59–61). Our data also suggest that confined ephrinB2 movement is correlated with reduced NiV/VSV pp infectivity. In summary, our data support a model in which IFITM1 sequesters ephrinB2 into GM1 membrane domains where its lateral mobility is restricted through a mechanism independent of membrane rigidity alone, thereby making receptor-mediated virus-cell membrane fusion less favorable and reducing the likelihood of productive virus-receptor encounters.

**Fig. 6.**
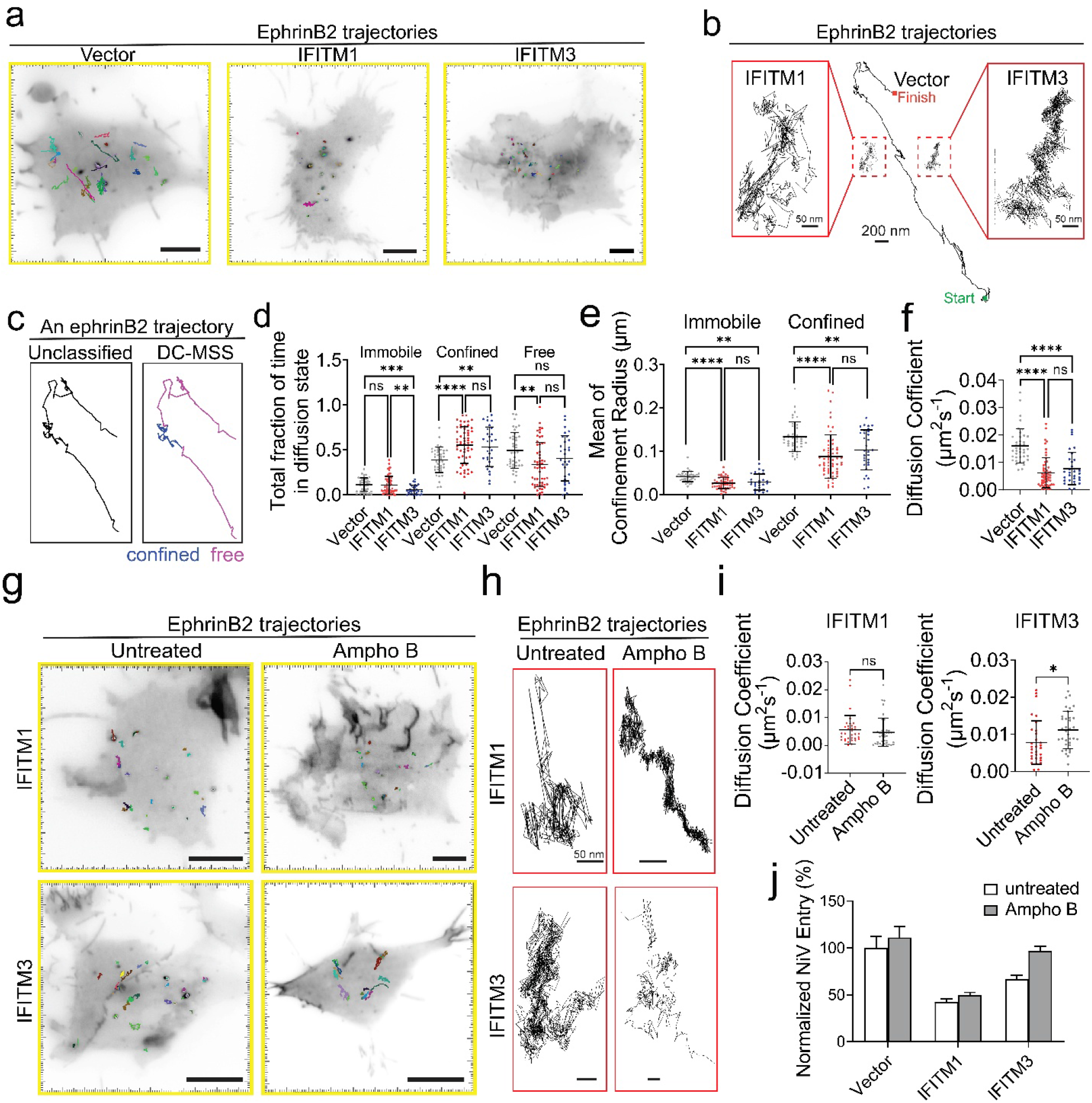
IFITM1 and IFITM3 restrict ephrinB2 lateral diffusion on the plasma membrane. **a**, ephrinB2 trajectories in HEK293T cells stably expressing IFITM1 (293T-IFITM1), IFITM3 (293T-IFITM3), and control cells. EphrinB2 was sparsely labeled by mStaygold (ephrinB2-RT-mStaygold) and tracked using TIRF microscopy at 30 ms/frame for 1 min. Scale bar: 5 μm. **b**, comparison of representative individual ephrinB2 trajectories in HEK293T-IFITM1, -IFITM3 cells and control cells. Trajectories are shown at the same scale (middle), and a zoomed-in view of an ephrinB2 trajectory from a 293T-IFITM1 (left) or 293T-IFITM3 (right) cell is provided. **c**, ephrinB2 trajectories were analyzed using DC-MSS, which segments trajectories by three different motion modes, immobile (black, not shown), confined (blue), and free (magenta). **d**, Mean fraction of time spent by ephrinB2 in each motion type in control cells (vector), 293T-IFITM1 (IFITM1), and 293T-IFITM3 (IFITM3) cells. Each dot represents the mean for one cell. **e** and **f**, Mean of confinement radius and diffusion coefficient (**f**) for the same trajectories as in (**d**). Sample size n = 40 (vector), 58 (IFITM1), and 31 (IFITM3) from at least three independent experiments. **g**, ephrinB2 trajectories in ampho B-treated and untreated 293T-IFITM1 cells (top) and 293T-IFITM3 cells (bottom). 293T-IFITM1 or -IFITM3 cells were transfected with ephrinB2-RT-mStaygold. At 24 hrs post-transfection, cells were treated with 1 μM ampho B or PBS for 1 hrs at 37°C. Then, Ampho B was removed and fresh medium was supplemented. Imaging was completed within 1 hr after the treatment. **h**, comparison of individual ephrinB2 trajectories in Ampho B-treated and untreated 293T-IFITM1 cells (top) and 293T-IFITM3 cells (bottom). Trajectories are shown at the same scale. **i**, mean of diffusion coefficient for the trajectories recorded from ampho B-treated and untreated 293T-IFITM1 and 293T-IFITM3 cells. Sample size: n = 34 (untreated, 293T-IFITM1 cells) and 41 (ampho B-treated, 293T-IFITM1 cells); 31(untreated, 293T-IFITM3 cells), and 35 (ampho B-treated, 293T-IFITM3 cells). **j**, NiV/VSV pp entry into 293T-IFITM1 and 293T-IFITM3 cells upon ampho B treatment. Control (vector), 293T-IFITM1, and 293T-IFITM3 cells were treated with 1 μM Ampho B (Ampho B, grey) or buffer (untreated, white) then infected with NiV/VSV pp. Virus entry was measured by luminescence intensity and normalized to untreated control. Bars represent means ± SEM. *p-*value was obtained using Welch’s *t*-test. (nonsignificant, ns; ***, *p* ≤ 0.001; ****, *p* ≤ 0.0001.). All experiments were repeated at least three times.

## Discussion

Our study reports that IFITM1 inhibits NiV and HeV glycoprotein-mediated virus entry and cell-cell fusion in both endothelial and epithelial cell lines. Using siRNA knockdown in endothelial cell lines (HuMEC and HuVEC), we identified that IFITM1 is the most potent IFITM protein in inhibiting the virus entry and cell-cell fusion mediated by HNV glycoproteins (Fig. 1 and Fig. 2). This agrees with a previous report showing that overexpression of IFITM1 prevented infection by paramyxoviruses, pneumoviruses, and herpesviruses, which enter cells at the plasma membrane (6). With IFN-α stimulation, endogenous IFITM2 significantly inhibits NiV/VSV pp and HeV/VSV pp in HEK293T cells (Fig. 1). Interestingly, IFITM3 knockdown only slightly increases virus entry in HEK293T cells with IFN-α stimulation. Due to the cross-reactivity of siRNAs targeting IFITM2 and IFITM3, the increase of the virus entry upon IFITM2 siRNA treatment may result from a synergistic knockdown of both IFITM2 and IFITM3 (Fig. S1). Overexpression of IFITM1 and IFITM3 in HEK293T and HeLa cells mostly restricts HNV entry and cell-cell fusion. It is believed that the antiviral activity of IFITM proteins is correlated with their subcellular localization. This localization is affected by the overall expression level and post-translational modification (4, 62). IFITM1 is constitutively expressed on the cell surface, thus is a more potent inhibitor of viruses that enter at the plasma membrane than IFITM3, which is mainly localized to endosomes and transiently present at the cell surface (Fig. 3). We noticed that overexpression of IFITM2 resulted in endosomal localization and significantly enhanced HNV entry to HeLa cells. Interestingly, IFITM3 is reported to promote NiV pseudovirus entry when expressed in the endosomes in MDCK cells (32). This is not surprising because NiV has been shown to enter cells via macropinocytosis (63). Additionally, endosomal IFITM2 can promote SARS-CoV-2 infection by interacting with the spike protein (12). These lines of evidence suggest that the antiviral effect of IFITMs is highly dependent on its sub-cellular localization.

The current view is that IFITM3 blocks the endosomal entry of IAV by increasing membrane rigidity (13, 23, 24) and modulating membrane curvature (14, 19). A recent study shows that IFITM3 locally redistributes membrane lipids, reducing cholesterol in its immediate vicinity and enriching lipids that disfavor fusion at the hemifusion diaphragm (26). This lipid sorting shifts the reaction toward nonproductive hemifusion intermediates and increases the energy barrier for fusion, which is manifested as arrest at hemifusion and inhibition of fusion pore enlargement (17, 23). This model places less emphasis on bulk membrane rigidity and instead highlights the nanoscale lipid composition and organization of the local membrane surrounding each fusion site. Complementing these findings, our SMLM data show that IFITM1 molecules form nanoclusters which co-enrich with the CtxB labeled-GM1 domain (Fig. 4 and 5). GM1 domains are often considered as raft-like domains with increased membrane rigidity than the bulk membrane and concentrate membrane receptors. EphrinB2 is a type I transmembrane ligand for the EphB3 receptor tyrosine kinase, and has been reported to be associated with lipid rafts, as shown by its enrichment in detergent-resistant, caveolin-associated membrane fractions (64). Accordingly, ephrinB2 clusters are also enlarged when correlated with IFITM1 (Fig. 5), potentially due to either direct IFITM1-ephrinB2 interactions, or co-targeting to GM1 membrane domains.

Although raft-like domains are considered energetically disfavored for membrane fusion, evidence from earlier studies shows that globally depleting membrane cholesterol or altering receptor partitioning to cholesterol-rich, raft-like membrane domains inhibits virus entry and attachment (65–68). It would have been plausible that IFITM1-mediated ephrinB2 sequestration in the cholesterol-dependent GM1 domain may favor NiV infection. Recent studies on modeled membranes show that although raft-like, liquid-ordered (L_o_) lipid domains are necessary and sufficient for HIV binding and membrane fusion, both processes occur preferentially at the boundary between L_o_ and liquid-disordered (L_d_) domains, where hydrophobic mismatch may help lower the energetic barrier to fusion (69–71). Thus, membrane fusion depends not only on the presence of cholesterol-dependent, L_o_ domains but also on local lipid-phase heterogeneity surrounding the receptor. In our system, IFITM1-associated ephrinB2 clusters are larger and denser (Fig. 5), and GM1 domains are also enlarged in the presence of IFITM1 (Fig. 5). Since ephrinB2 clustering is thought to promote NiV-G engagement (72), IFITM1 may simultaneously redistribute a greater fraction of ephrinB2 into enlarged raft-like membrane domains, presumably reducing the relative extent of L_o_/L_d_ interfaces that may be needed to support efficient membrane fusion.

Current evidence suggests that productive virus entry requires a balance between receptor mobility and confinement. Sufficient receptor mobility can facilitate virus-receptor encounter and recruitment of receptors to virus attachment sites (73, 74), whereas local confinement after engagement can help stabilize the fusion machinery at the virus-cell interface (72, 75–77). Our data show that the mobility of ephrinB2 receptor is constrained in 293T-IFITM1 cells. This may lead to reduced virus entry by two mechanisms: 1) the reduced lateral diffusion of ephrinB2 receptor may lower the probability of virion-receptor encounter and limit receptor recruitment to the virus attachment site, thereby reducing virus attachment; 2) even when virions engage ephrinB2 molecules, the confinement of ephrinB2 may still permit the assembly of fusion machinery, however, productive membrane fusion events may remain inefficient because IFITM1 is expected to make the local membrane environment less permissive for fusion via increasing membrane rigidity and/or reducing the phase heterogeneity surrounding ephrinB2. Larger and denser IFITM1-associated ephrinB2 clusters could contribute to reduced receptor mobility, as larger assemblies would be expected to diffuse more slowly. However, this mechanism alone is unlikely to explain the overall shift in ephrinB2 dynamics, because most ephrinB2 tracks in 293T-IFITM1 cells are confined and less diffusive (Fig. 6a). In contrast, only a subset of ephrinB2 clusters is detectably associated with IFITM1 (Fig. 5f). These observations suggest that direct clustering with IFITM1 accounts for only part of the constrained ephrinB2 mobility, and that IFITM1 must also influence a broader membrane environment that restricts ephrinB2 diffusion beyond the receptors present in obvious IFITM1-associated clusters. We attempted to rescue ephrinB2 lateral diffusion by using ampho B, based on the hypothesis that reduced ephrinB2 mobility in IFITM1-expressing cells is caused by increased membrane rigidity and could be rescued by the membrane-permeabilizing agent, ampho B (24). However, ampho B did not have an impact on either ephrinB2 diffusion nor NiV/VSV pp entry in 293T-IFITM1 cells (Fig.6 g-j), although it did counteract IFITM3-mediated NiV restriction and enhanced ephrinB2 diffusion in 293T-IFITM3 cells (Fig. 6 g-j). This is not entirely unexpected, because ampho B was previously shown to relieve IFITM2- and IFITM3-mediated restriction, but not IFITM1-mediated restriction, in studies that used influenza A virus which enters via endocytosis. These data suggest that IFITM1 confines ephrinB2 through mechanisms other than co-clustering and modulating bulk membrane rigidity in an ampho B-reversible manner. One possibility is that IFITM1 alters ephrinB2 interactions with the cortical actin network, or the nanoscale organization of cortical actin itself, thereby restricting receptor diffusion (78, 79).

In summary, our study shows that IFITM proteins can inhibit HNV entry and cell-cell fusion in epithelial and endothelial cells by sequestering the ephrinB2 receptor in a fusion-unfavorable membrane domain, thereby increasing the barrier for virus-cell membrane fusion and decreasing the likelihood of virus-receptor encounter.

## Materials and Methods

### Plasmids

FLAG-tagged IFITM1, IFITM2, and IFITM3 genes were ligated into a pQCXIP (Takara, 631516) expression plasmid flanked by BamHI and EcoRI restriction sites, respectively. The VSV-G expressing plasmid, pMD2.G, was a gift from Didier Trono (Addgene plasmid # 12259). PcDNA3.1-V5-tagged ephrinB2 plasmid was a gift from Benhur Lee at Mount Sinai School of Medicine (30). PcDNA3/F-tractin-mStayGold was a gift from Atsushi Miyawaki (Addgene plasmid # 212019) (80). The gene encoding V5-tagged ephrinB2 was fused to a translational readthrough (RT) sequence (CAATAGGGCTTA) at its C-terminus, enabling sparse labeling, followed by the fluorescent protein mStayGold (58, 81). The construct was subsequently cloned into the pcDNA3.1 vector. Plasmids coding for HA-tagged NiV-G and FLAG-tagged NiV-F were described previously (47). HeV-F (NP_047111.2) and G (AAV80426.1) genes were synthesized by Geneart and modified as described previously (82). A FLAG-tag was inserted between amino acid residue 104L and 105V in HeV-F. An HA tag was added to the C-terminus of HeV-G. Both HeV-F and G constructs were cloned into pcDNA3.1 vector.

### Cell lines

HeLa cells, PK13 cells, and Human embryonic kidney (HEK) 293T cells were cultured in Dulbecco’s modified Eagle’s medium (DMEM, Sigma-Aldrich D6429) with 10% fetal bovine serum (FBS, Thermo Fisher 12483-020). Human microvascular endothelial cells (HuMEC) (ATCC CRL-4060) were cultured in Vascular cell basal medium (ATCC, PCS-100-030) with Microvascular endothelial cell growth kit – BBE (ATCC, PCS-110-040) and 0.5 ug/ml puromycin (10 mg/ml stock, Gibco A1138-03). Cells were washed with phosphate-buffered saline (PBS, Thermo Fisher Scientific, 10010049) and detached by 0.25% trypsin-EDTA (Thermo Fisher Scientific, 25200072) before seeding and sub-culturing. Trypsin-EDTA for primary cells (Sigma-Aldrich, T3924-100 ml) and Trypsin inhibitor from soybean (Sigma-Aldrich, T6414-100 ml) were used for HuMEC cells passaging and seeding. Human Umbilical Vein Endothelial Cells (HuVEC) (PromoCell, C-12203) were cultured in endothelial cell growth medium, which consists of SupplementMix (PromoCell, C-39215) and endothelial cell growth medium (PromoCell, C-22010B). HuVEC were treated with DetachKit-30 consisting of Hepes BSS (30 mM) (PromoCell, C-40000), Trypsin/EDTA (PromoCell, C-41000), and TNS (PromoCell, C-41100) before detaching and subculturing. To generate stable cell lines, retroviral pQCXIP vectors encoding FLAG-tagged IFITM constructs (6 μg), MLV-Gag/Pol (6 μg), and pmd2.G (3 μg) plasmids were co-transfected using polyethylenimine (PEI) at 1 mg/ml (Polysciences, 23966-100) to HEK293T cells cultured in a 10-cm dish. Supernatant was collected 48 hrs post-transfection to infect HEK293T cells or HeLa cells. After 3 days of infection, 2 μg /ml puromycin selection was applied for 28 days to establish a stable cell line expressing the FLAG-tagged IFITM constructs.

### siRNA, antibodies, and drugs

The siRNAs used in this study are: non-targeting siRNA (Horizon Discovery, D-001206-13-05), IFITM1 (Horizon Discovery, M-019543-01-0005), IFITM2 (Horizon Discovery, M-020103-02-0005) and IFITM3 (Horizon Discovery, M-014116-01-0005) specific siRNAs. Antibodies used in this study are: anti-FLAG mouse monoclonal antibody (Sigma-Aldrich, F1804); purified anti-HA.11 epitope tag antibody (Biolegend, 902301); anti V5-Tag (D3H8Q) rabbit monoclonal antibody (Cell Signaling Technology, 13202S); anti-GAPDH mouse monoclonal antibody (6C5) (Sigma, CB1001); IFITM1-specific monoclonal antibody (Proteintech, 600714-IG); IFITM2 polyclonal antibody (Proteintech, 12769-1-AP); IFITM3 polyclonal antibody (Proteintech, 11714-1-AP); human/mouse/rat ephrinB2 antibody (R&D systems, AF496); recombinant human EphB3-Fc chimera protein (R&D systems, 5667-B3); anti-mouse goat polyclonal antibody, HRP conjugated (Bio-Rad, 1705047); and anti-rabbit goat polyclonal antibody, HRP conjugated (Bio-Rad, 1706515); anti-mouse donkey polyclonal antibody, Alexa Fluor 647 conjugated (Invitrogen, A31571); anti-goat donkey polyclonal antibody, Alexa Fluor 647 (Invitrogen, A21447); Cholera Toxin Subunit B (CtxB), Alexa Fluor™ 647 conjugated (Invitrogen, C34778); anti-mouse donkey polyclonal antibody, Alexa Fluor 488 conjugated (Invitrogen, A21202); anti-human donkey polyclonal antibody, Alexa Fluor 488 conjugated (Jackson ImmunoResearch, 709-545-098); The cy3B fluorophore (Cytiva, PA63101) is conjugated to the donkey anti-mouse antibody (Jackson ImmunoResearch, 705-001-003) in-house by Ablabs. Amphotericin B (Ampho B; Sigma-Aldrich, A2942).

### Pseudovirus production

HEK293T cells were seeded in poly-D-lysine-coated 15 cm cell culture dish before transfection. At a 70%-80% confluency, pcDNA3.1 plasmids coding for FLAG-tagged F and HA-tagged G were transfected at a 3:1 ratio for NiV or 1:3 ratio for HeV using PEI. The pCDNA3.1 vector was transfected as the negative control. At 14-16 hrs post-transfection, cells were infected with recombinant VSV viruses, in which the gene coding for VSV-G was replaced by a Renilla luciferase gene, diluted in infection buffer (0.1% fetal bovine serum in PBS). At 2 hrs post-infection, infection buffer was replaced by pre-warmed complete growth medium, cells were incubated at 37°C 5% CO_2_ incubator for 48 hrs. The supernatant containing NiV/VSV and HeV/VSV pseudoviruses were collected, filtered by 0.22 μm filters to remove cell debris, and concentrated by ultra-centrifugation through a 20% sucrose cushion in NTE buffer (150 mM NaCl, 40 mM Tris-HCl, 1 mM EDTA, pH 8.0) at 26,400 rpm (100,000 × g) at 4°C for 1.5 hrs. The supernatant was removed, and the pellet was resuspended in the appropriate amount of 5% sucrose in NTE buffer. Aliquots were stored at -80°C for use. The expression levels of NiV-F and -G proteins in pseudoviruses were verified by Western Blot, and the VSV genome copy number was quantified by quantitative RT-PCR as previously described (83).

### Western Blot analysis

HEK293T or HeLa cells stably expressing IFITM proteins or HEK293T, HuMEC, and HuVEC transfected with IFITM-specific siRNAs were seeded in 12-well plates. At 90% confluency, 100 μl 1× RIPA buffer (Sigma-Aldrich, 20-188) with protease inhibitor (Sigma-Aldrich, 11836170001) was added into each well, incubated on ice for 20 min, and then centrifuged at 4°C, 13,000 rpm (16,000 × g) for 20 min. The supernatant was collected and mixed with 1× SDS-PAGE sample loading buffer (60 mM Tris-HCl (pH=6.8); 2% SDS; 10% glycerol, 0.025% Brophenol blue) supplemented with 15 mM dithiothreitol (DTT; Thermo Fisher, R0861). All samples were heated at 95°C for denaturation. The denatured samples were loaded into 10% polyacrylamide gels for SDS-PAGE. Proteins were transferred to a 0.45 μm PVDF membrane (Cytiva, GE10600021). The PVDF membranes were blocked with 1% bovine serum albumin (BSA; Sigma-Aldrich, A9647) in PBS for 60 min at room temperature, followed by incubation with specific primary and secondary antibodies. Protein bands were visualized using Clarity Western ECL substrate (BioRad, 1705060) and captured with a ChemiDoc MP Imaging System (BioRad).

### Coimmunoprecipitation

HEK293T cells seeded in a 6-well plate were transfected with the following plasmid combinations: 1) 2.5 μg empty pcDNA 3.1 vector, 2) 1.25 μg empty pcDNA 3.1 vector and 1.25 μg FLAG-tagged IFITM1, 3) 1.25 μg empty pcDNA 3.1 vector and 1.25 μg V5-tagged ephrinB2, 4) 1.25 μg FLAG-tagged IFITM1 and 1.25 μg V5-tagged ephrinB2 using Lipofectamine 3000. Each sample is duplicated. At 48 hrs post-transfection, cells from 1 well of a 6-well plate were washed with pre-warmed PBS and lysed in 200 μl low-salt lysis buffer (1% EcoSurf EH-9, 50 mM Tris HCl (pH 8.0)) and supplemented with protease inhibitors. Cells were incubated on ice for 30 min, followed by centrifugation at 16,000 x g for 20 min at 4°C. 60 μl of cell lysate was set aside for immunoblot analysis, and the rest was used for immunoprecipitation, as recommended by the manufacturer. 6 μl anti-DYKDDDDK microbeads (Miltenyi Biotec, 130-101-591) were added to 140 μl cell lysates and incubated for 30 min on ice. μ columns (Miltenyi Biotec, 130-101-591) were prepared according to the manufacturer’s instructions. Lysate was run over the columns, and microbeads were washed according to the manufacturer’s instructions. 20 μl preheated elution buffer (95°C) was added to the column before eluting the bound immunoprecipitated protein in 50 μl elution buffer. Elutes were resolved by 10% SDS-PAGE, and proteins were immunoblotted with mouse anti-FLAG, rabbit anti-V5, and mouse anti-GAPDH primary antibodies, followed by HRP-conjugated goat anti-mouse and goat anti-rabbit secondary antibodies. Protein bands were visualized as described above.

### Cell-cell fusion assay

For the homologous cell-cell fusion assay, HEK293T or HeLa cells stably expressing IFITM proteins were seeded in 12-well plates before transfection. At 70%-80% confluency, 1 μg total DNA containing FLAG-tagged NiV-F and HA-tagged NiV-G at a 3:1 ratio or FLAG-tagged HeV-F and HA-tagged HeV-G at a 1:3 ratio was transfected using Lipofectamine 3000 (Thermo Fisher, L3000015). At 18-24 hrs post-transfection, cells were fixed by 4% PFA. Cell-cell fusion was observed using a Nikon TE2000U microscope at a 10× magnification. Images of five different fields in each well were taken for syncytia counting. For the heterologous cell-cell fusion assay, HEK293T cells or HuVEC (target cells) were seeded in a 12-well plate at 70%-80% confluency, and then transfected by siRNA targeting IFITM proteins. PK13 cells (effector cells) were seeded in 10 cm plates. At 70%-80% confluency, 10 ug total DNA containing FLAG-tagged NiV-F and HA-tagged NiV-G at a 3:1 ratio or FLAG-tagged HeV-F and HA-tagged HeV-G at a 1:3 ratio was transfected using Lipofectamine 3000. After 24 hrs, transfected PK13 cells were detached and overlaid on HEK293T cells or HuVEC at a 1:3 or 1:1 ratio, respectively. After 2-3 hrs for HEK293T cells or 6-8 hrs for HuVEC, cell-cell fusion was observed and quantified as described above.

### siRNA knockdown

HEK293T cells were transfected with 40 pmol/well in a 12-well plate or 4 pmol/well in a 96-well plate of scrambled siRNA or IFITM specific siRNAs using RNAiMAX (Thermo Fisher, 13778100) at a 1:3 ratio in Opti-MEM (Gibco, 31985047). Twenty-four hours after the initial transfection, HEK293T cells were subjected to a second round of siRNA transfection. At 6-8 hrs after the second transfection,1000 IU Gibco IFN-α2b was added into HEK293T cells to stimulate IFITMs expression. At 16 hrs post-treatment, medium was removed, cells were washed once with pre-warmed PBS and subjected to further analysis. HuMEC were transfected with 20 pmol/well siRNA in a 12-well plate or 2 pmol/well siRNA in a 96-well plate as described above. At 24 hrs post-transfection, 100 IU Gibco IFN-α2b were added into HuMEC to stimulate IFITMs expression. At 16 hrs post-treatment, medium was removed, cells were washed once with pre-warmed PBS, and subjected for further analyses. HuVEC cells were transfected with 20 pmol/well siRNA in a 12-well plate as described above. At 24 hrs post-transfection, 1000 IU Gibco IFN-α2b was added into HuVEC cells to stimulate IFITMs expression. At 16 hrs post-treatment, medium was removed, cells were washed once with pre-warmed PBS and subjected to further analyses.

### qPCR analysis

The total RNA of HEK293T cells, HuMEC, and HuVEC transfected with scrambled and IFITM-targeting siRNAs was extracted by RNeasy Plus Mini Kit (Qiagen, 74134) following the manufacturer’s protocols. Extracted total RNAs were reverse transcribed using the SuperScript III first-strand synthesis system (Invitrogen, 18080051). mRNA expression levels were detected by IFITM-specific primers and probes. GAPDH is the internal control. Oligonucleotides used in qPCR are listed in Table 1.

**Table 1.**
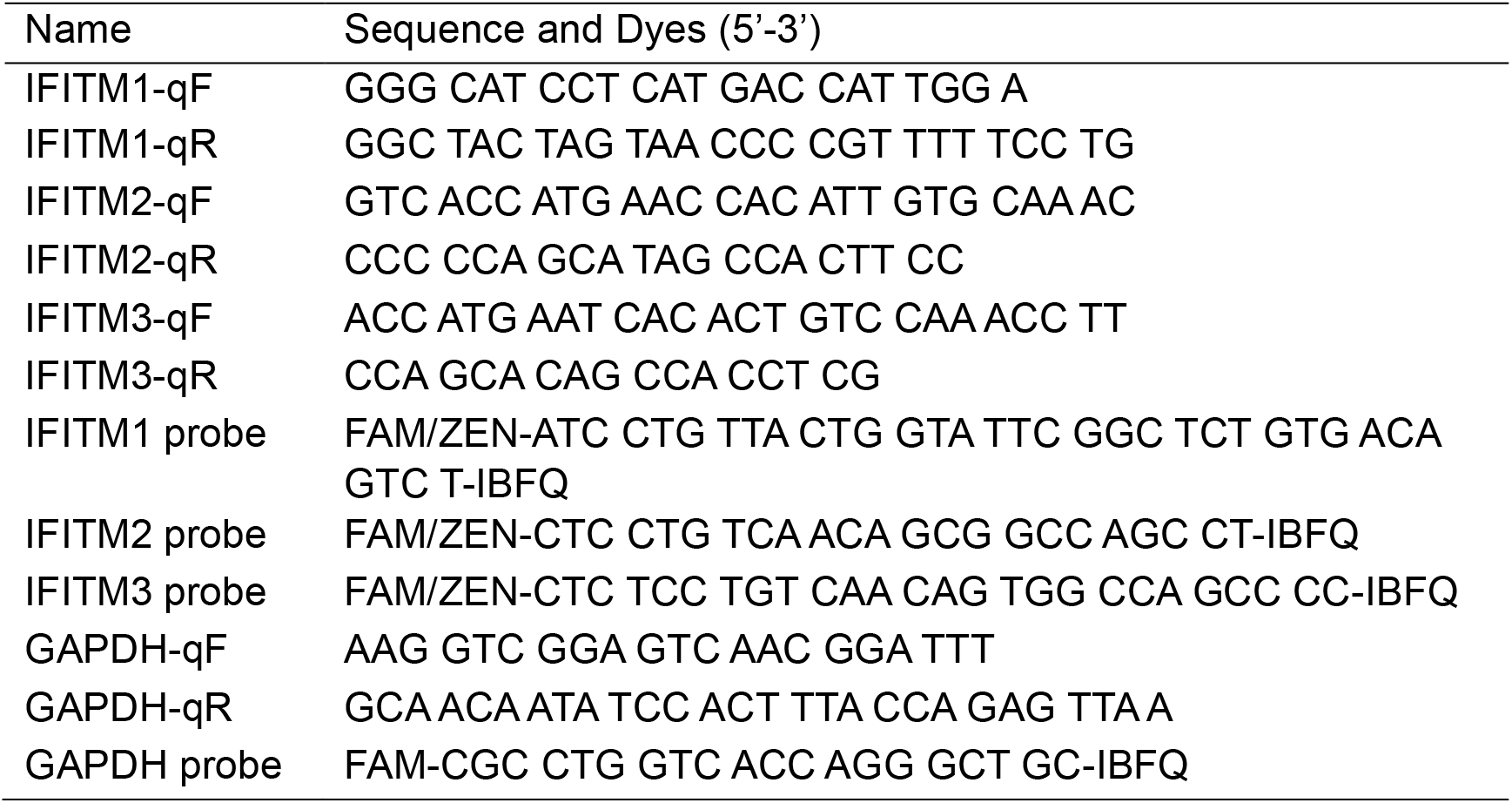
Oligonucleotides used in qPCR.

### Immunofluorescence for lattice SIM

4×10^5^ HEK293T or 1×10^5^ HeLa cells stably expressing IFITM proteins were seeded on 12 mm coverslips (Thermo Fisher, 1254580) coated with 1.25 μg of fibronectin (Sigma-Aldrich, F4759-2mg) in a 24-well plate. After 24 hrs, cells were washed by pre-warmed PBS and fixed by 4% paraformaldehyde (PFA) for 30 minutes at room temperature. After fixation, cells were briefly washed with PBS and permeabilized by 0.1% Triton X-100 at room temperature for 5 min. Cells were blocked with BlockAid (Life Technologies, B10710) for 1 hr at room temperature. The FLAG-tagged IFITM proteins were detected by the anti-FLAG mouse monoclonal antibody (Sigma Aldrich, F1804) and Alexa Fluor 488-conjugated, donkey anti-mouse polyclonal antibody (Invitrogen, A21202). Cells were incubated with primary antibody overnight at 4°C, and then with the secondary antibody for 1 hr at room temperature. Each antibody incubation was followed by five PBS washes, 5 min each time. Coverslips were embedded on glass slides with Prolong™ Diamond Antifade Mountant (Thermo Scientific, P36961). Fluorescence images were acquired using a Zeiss Elyra 7.

### Virus entry assay

4×10^4^ HEK293T, 1×10^4^ HeLa cells stably expressing IFITM proteins were seeded in a 96-well plate coated with Poly-D-lysine before infection. Alternatively, 4×10^4^ HEK293T cells and 1×10^4^ HuMEC were seeded and transfected by siRNAs as described above. After 24 hrs, the HNV/VSV pseudovirus was diluted in infection buffer and added to each well. At 2 hrs postinfection, the infection buffer was replaced by pre-warmed complete growth medium, and the plate was incubated at 37°C, 5% CO_2_ incubator. After 24 hrs, cells were lysed by 100 μl luciferase lysis buffer (Promega, E2810). The luciferase intensity was measured by using a Promega Renilla luciferase assay kit (Promega, E2810) on a TECAN Spark 10 M fluorescent plate reader.

### Cell viability assay

Cell viability was assessed using the Cell Counting Kit-8 (CCK-8; Abcam, ab228554) according to the manufacturer’s instructions. 8×10^4^ HEK293T cells or 2×10^4^ HuVEC were seeded in 96-well plates and incubated overnight at 37°C, 5% CO_2_. On the following day, cells were transfected with siRNAs as described above. The medium was replaced with 100 μl of fresh culture medium, and 10 μl of WST-8 reagent was added to each well. Plates were incubated at 37°C for 1 h. Fluorescence was measured at 460 nm using a TECAN Spark 10 M fluorescent plate reader to quantify cell viability.

### Flow cytometry

HEK293T cells transfected with IFITMs-targeting siRNAs or stably expressing IFITM proteins were collected for flow cytometry analyses. Briefly, cells were washed twice with pre-warmed PBS and detached from the plate using 1 ml phosphate-buffered saline (PBS) containing 10 mM EDTA. The cell suspension was centrifuged at 300×g for 5 min at 4°C. The cell pellets were fixed and permeabilized by using BD Cytofix/Cytoperm™ solution (BD Biosciences, 554722), incubated on ice for 20 min followed by two washes using BD Perm/Wash™ Perm/Wash Buffer (BD Biosciences, 554723). Cells were incubated with 10 μg/ml recombinant human EphB3 Fc Chimera Protein (R&D Systems, 5667-B3-050) on ice for 1 hr. After two washes by BD Perm/Wash™ Perm/Wash Buffer, cells were incubated with Alexa 488-conjugated, anti-human donkey polyclonal antibody (Jackson ImmunoResearch, 709-545-098) at a 1:400 dilution on ice for 45 min. Following two final washes, samples were analyzed by flow cytometry (Attune® NxT™ Acoustic Focusing Cytometer, Thermo Fisher). The results were analyzed using FlowJo. The mean fluorescent intensities (MFI) were normalized to the MFIs of the scrambled siRNA-transfected cells (for siRNA knockdowns) or empty vector control (for stable cells).

### Immunofluorescence for SMLM

For single-color SMLM imaging of IFITM1 in HEK293T cells, 4×10^5^ HEK293T cells were seeded on coverslips (Marienfeld, no. 1.5H, 18 mm) coated with 2.5 μg of fibronectin (Sigma-Aldrich, F4759-2mg) in a 12-well plate and allowed to adhere overnight at 37°C. On the following day, 1000 IU Gibco IFN-α2b was added into HEK293T cells for 16 hrs to stimulate IFITMs expression. For single-color SMLM imaging of ephrinB2 in HEK293T cells, cells were transfected with 1 μg V5-tagged ephrin B2 plasmid using Lipofectamine 3000. After IFN-α2b treatment (IFITM1 imaging) or 24 hrs post-transfection (ephrinB2 imaging), cells were fixed with PBS containing 4% paraformaldehyde (PFA; Electron Microscopy Sciences; 50980487) and 0.2% glutaraldehyde (Sigma-Aldrich, G5882-50 ml) for 90 min at room temperature. After fixation, cells were briefly washed with PBS and permeabilized by 0.1% Triton X-100 at room temperature for 5 min. Cells were treated with signal enhancer image-IT-Fx (Life Technologies, I36933) for 30 min at room temperature, and then blocked using BlockAid (Life Technologies, B10710) for 1 hr at room temperature. The endogenous IFITM1 proteins were detected by IFITM1-specific monoclonal antibody (Proteintech, 600714-IG) and an Alexa Fluor® 647 conjugated donkey anti-mouse secondary antibody (Invitrogen, A31571). Ephrin B2 proteins were detected by human/mouse/rat ephrinB2 antibody (R&D systems, AF496) and anti-goat donkey polyclonal antibody, Alexa Fluor 647 (Invitrogen, A21447). Cells were incubated with primary antibody overnight at 4°C, and then with the secondary antibody for 1 hr at room temperature. Each antibody incubation is followed by five PBS washes, 5 min each time. Cells were then fixed in PBS containing 4% PFA for 10 min at room temperature.

For dual-color SMLM imaging to visualize the spatial distribution of ephrinB2 and IFITM1, HEK293T cells were seeded, incubated as described above, and then transfected with 0.5 μg V5-tagged ephrin B2 plasmid and 0.5 μg FLAG-tagged IFITM1 plasmid. Cells were fixed and permeabilized as described above. Ephrin B2 proteins were detected by human/mouse/rat ephrinB2 antibody (R&D systems, AF496) and anti-goat donkey polyclonal antibody, Alexa Fluor 647 (Invitrogen, A21447). For dual-color SMLM imaging to visualize the distribution of GM1 domain and IFITM1, HEK293T cells were seeded in a 12-well plate and transfected with 1 μg FLAG-tagged IFITM1 plasmid using Lipofectamine 3000. At 24 hrs post-transfection, cells were fixed with 4% paraformaldehyde and 0.2% glutaraldehyde at room temperature for 90 min. Cells were incubated with an Alexa Fluor® 647 conjugated Cholera Toxin Subunit B (Invitrogen, C34778) to detect GM1, followed by a secondary fixation using 4% paraformaldehyde and 0.2% glutaraldehyde, and permeabilization with 0.1% Triton×100. The FLAG-tagged IFITM1 proteins were detected by the anti-FLAG mouse monoclonal antibody (Sigma-Aldrich, F1804) and a Cy3B-conjugated donkey anti-mouse secondary antibody.

### SMLM setup, imaging, and data analysis

SMLM was performed on a custom-built microscopy described previously (46, 47, 84) . Fluorescence beads (Life Technologies, F8799) were added to samples as fiducial markers for drift control. Samples were immersed in imaging buffer as described previously (85). For SMLM imaging, samples were exposed to a laser power density of 1 kW/cm^2^ for the 639- and 532-nm lasers to activate Alexa Fluor 647 and cy3B, respectively. A total of 40,000 images were acquired at 50 Hz to reconstruct one SMLM image. Custom-written software in MATLAB (MathWorks) was used to reconstruct SMLM images. Clusters of NiV-M localizations were identified and characterized using ClusDoC (48). The min points and ε for DBSCAN were set at 3 and 10, respectively. The degree of colocalization (DoC) assay was performed using ClusDoc. Each localization in channel 1 is assigned a degree of colocalization (DoC) value according to its correlation with the localization in channel 2. The DoC value ranges from -1 to 1, with -1 indicating anti-correlation and 1 correlation. A localization in channel 1 must have a DoC value > 0.4 to be considered as correlated with a localization in channel 2. A colocalized cluster must contain more than 10 localizations with DoC values > 0.4 (48, 86).

### TIRF microscopy and single particle tracking (SPT)

For SPT experiments, 3.2 ×10^5^ HEK293T cells stably transduced with pQCXIP-IFITM1 (293T-IFITM1) or empty vector were seeded in a glass bottom 35 mm Dish (Ibidi, 81218-200) coated with 5 μg of fibronectin (Sigma-Aldrich, F4759-2mg) and allowed to adhere overnight at 37°C. On the following day, cells were transfected with pcDNA3.1 plasmid expressing an ephrinB2 construct fused with a C-terminal RT sequence, followed by an mStayGold fluorescent tag (ephrinB2-RT-mStaygold). At 20-24 hrs post transfection, cells were subjected to TIRF imaging. In the case of Ampho B treatment, 1 μM Ampho B (Sigma-Aldrich, A2942) or PBS was added to the growth medium of 293T-IFITM1 transfected with ephrinB2-RT-mStaygold at 24 hrs post transfection. Cells were incubated at 37°C, 5% CO_2_ for 1 hr. Then, cells were replenished with fresh pre-warmed complete growth medium and subjected to TIRF imaging for 1 hr. TIRF Imaging was performed by a Zeiss Elyra7 microscope equipped with a 63× NA 1.46 objective lens at TIRF illumination. A live cell chamber maintained a 37°C and 5% CO_2_ atmosphere. Images were acquired at 30 ms per frame for 1 min. The image analysis was performed in 6 steps by Imaris (Oxford Instruments): (i) particle detection; (ii) particle properties (e.g. positions and intensities) on each frame; (iii) frame-to-frame particle tracking; (iv) gap closing; (v) complete track identification; (vi) track filtering by track duration. Tracks that lasted over 5 s were included in further analysis. Tracks were analyzed using DC-MSS (divide-and-conquer moment scaling spectrum), which combines sliding-window track segmentation with moment scaling analysis to resolve transient mobility states along individual tracks (87). Briefly, a local sliding-window moves throughout a track to partition it into segments of putatively different motion classes. These segments are classified into distinct motion states via moment scaling spectrum analysis of molecule displacements, including immobile, confined, free diffusion. Track segmentation is refined by combining short segments with adjacent classified segments to achieve a classification of the original or a lower mobility class, according to the MSS analysis results. DC-MSS enables the detection of dynamic state transitions within each trajectory, thereby capturing heterogeneous and short-lived motions that would otherwise be obscured by ensemble averaging. The confinement areas of immobile and confined motions, the diffusion coefficients, and the time fraction spent in each motion type were computed on a segment-by-segment basis (87).

### Statistical analysis

Statistical analyses were conducted using GraphPad Prism 9.0. Specific statistical tests, sample sizes (n), and the number of independent biological replicates are detailed in the respective figure legends.

## Data, materials, and software availability

All data is included in the article. Materials are available upon request to Dr. Qian Liu.

## Acknowledgements

This work was supported by funding from the Canadian Institutes of Health Research (CIHR; Project Grant, PJT-183861) and McGill Interdisciplinary Initiative in Infection and Immunity (MI4)-Pfizer Early Career Investigator Award to Q.L; CIHR Project Grant PJT-166048 to C.L.. Y.L. and Q.W. are supported by PhD fellowships from Fonds de recherche du Québec – Nature et technologies (FRQNT) and Chinese Scholarship Council. J.W. is supported by a postdoctoral fellowship from FRQNT. We thank Dr. Youssef Chebli at the Multi-Scale Imaging Facility Services at McGill University for technical support. Q.L. thanks the late Dr. Keng Chou from the Department of Chemistry at the University of British Columbia for his mentorship, encouragement, and lasting support throughout her scientific development.

## Author contributions

Y.L., C.L., Q.L. designed research; all authors performed research; J.L. and Q.L. developed data analysis software and codes; Y.L., J.W., J.L., and Q.L. analyzed data; Y.L. and Q.L. wrote the paper with edits from all authors. C.L. and Q.L. supervised research and acquired funding.

